# High resilience of the mycorrhizal community to prescribed seasonal burnings in a Mediterranean woodland

**DOI:** 10.1101/2020.06.10.141671

**Authors:** Stav Livne-Luzon, Hagai Shemesh, Yagil Osem, Yohay Carmel, Hen Migael, Yael Avidan, Anat Tsafrir, Sydney I. Glassman, Thomas D. Bruns, Ofer Ovadia

## Abstract

Fire effects on ecosystems range from destruction of aboveground vegetation to direct and indirect effects on belowground microorganisms. Although variation in such effects is expected to be related to fire severity, another potentially important and poorly understood factor is the effects of fire seasonality on soil microorganisms. We carried out a large-scale field experiment examining the effects of spring versus autumn burns on the community composition of soil fungi in a typical Mediterranean woodland. Although the intensity and severity of our prescribed burns were largely consistent between the two burning seasons, we detected differential fire season effects on the composition of the soil fungal community, driven by changes in the saprotrophic fungal guild. The community composition of ectomycorrhizal fungi, assayed both in pine seedling bioassays and from soil sequencing, appeared to be resilient to the variation inflicted by seasonal fires. Since changes in the soil saprotrophic fungal community can directly influence carbon emission and decomposition rates, we suggest that regardless of their intensity and severity, seasonal fires may cause changes in ecosystem functioning.

**Declarations:** *Funding:* This research was co-supported by the United States-Israel Binational Science Foundation (BSF Grant 2012081) and Tel-Hai College.

*Conflicts of interest/Competing interests:* We declare no conflicts of interest and that this material has not been submitted for publication elsewhere.

*Ethics approval:* Not applicable

*Consent to participate:* Not applicable

*Consent for publication:* Not applicable

*Availability of data and material:* Sequences were submitted to the National Center for Biotechnology Information Sequence Read Archive under accession numbers SRRXXX◻SRRXXX.

*Code availability:* Not applicable

*Authors’ contributions:* OO HS TB YO YC conceived and designed the experiment. SSL YA HM AT performed the experiment. SIG provided the pipeline scripts, and guidance in bioinformatics work and analyses. SLL OO HS wrote the paper and analyzed the data, and all authors contributed substantially to revisions.

## Introduction

Fire is one of the most common natural and anthropogenic disturbances leading to secondary succession of both plant and fungal communities (Marlon et al. 2009). Exploring the effects of fire on ecosystem functioning is of high priority, especially due to the increase in fire risk associated with climate change (Moriondo et al. 2006; Pechony and Shindell 2010; Westerling et al. 2006). The extent of damage fires inflict on plant communities is manifested not only through the destruction of plant tissues, but also in destruction of symbiotic soil microbes, which may be necessary to buffer against fire effects, thus increasing plant community resilience (Johnstone et al. 2010; Kipfer et al. 2011). Most of the temperate and boreal trees around the globe are obligately symbiotic with ectomycorrhizal fungi (EMF), meaning that their establishment is dependent upon the occurrence of an appropriate symbiont community (Miller et al. 1998). Therefore, fire effects on the belowground biota may be far-reaching with regard to vegetation regeneration and growth during the first few post-fire years (Neary et al. 1999). For example, a fire study based on chrono-sequence found that fire temporarily shifted the fungal community structure and function by increasing the abundance of saprotrophic fungi (Sun et al. 2015). Eventually the community returned to its pre-fire state, but at a very slow rate (Sun et al. 2015). Such a community shift towards saprotrophic fungi may have a detrimental effect on ecosystem functioning because it may shift the balance between obligate symbiotic EMF, associated with tree roots, and saprotrophic fungi. Besides the clear negative outcome of reduced symbionts available for plants (Collier and Bidartondo 2009), competition between these two fungal guilds can suppress decomposition rates (i.e., the Gadgil effect, Fernandez and Kennedy 2016; Gadgil and Gadgil 1975; Gadgil and Gadgil 1971).

Various studies have demonstrated both direct and indirect effects of fire on the EMF community while consequently influencing the post-fire regeneration of the plant community (Buscardo et al. 2010; Glassman et al. 2016b; Johnson 1995; Marlon et al. 2009; Miller and Urban 1999; Taudière et al. 2017; Veen et al. 2008). Although such effects are expected to be related to fire severity, which often varies during the year, less is known about the specific effect of fire season on the EMF community. The aboveground importance of fire season is well established. Specifically, compared with spring fires, autumn fires consume greater portions of the landscape area, standing plant biomass and other organic material (Knapp et al. 2005), while having more profound negative effects on the understory vegetation richness (see Knapp et al. 2009 for a thorough review). However, less attention has been given to the effect of fire season on the subterranean part of the ecosystem (but see, de Roman and de Miguel 2005; Smith et al. 2004).

Examining the effects of prescribed burns on the EMF community in a natural setting of ponderosa pine stands in eastern Oregon, Smith *et al.* (2004) found that autumn fires had long lasting, devastating effects on the mycorrhizal community, with a reduction of 80% in molecular species richness. Spring fires, however, did not differ from the unburned control. Smith et al. (2004) suggested that observed differences in the EMF community composition were the result of inter-season variation in fire severity. Specifically, the low moisture content in the fuel and in the soil during late season, resulted in higher soil temperatures and increased microbial mortality. Such extreme soil temperatures may damage the mycorrhizal community directly by destroying the mycelia, or indirectly by host death, both can lead to a long lasting negative effect on the EMF community (Klopatek et al. 1994). On the contrary, spring fires usually occur after the wet season when soil moisture is high and heat transfer is highly efficient, compensating for the increase in soil temperature caused by these fires, and thus having a weaker detrimental effect on the soil biota. Seasonal fire effects can be also related the phenological stages of both plants and fungi, resulting in a differential effect on their community composition. For example, in many fungal species characterizing Mediterranean habitats, the amount of mycelium decreases in summer, probably due to hot and dry conditions, whereas in autumn it increases again (De la Varga et al. 2013). Furthermore, during the hot dry Mediterranean summer selection may favor fungal species which can better cope with these extreme conditions, resulting in a seasonal shift in the composition of the fungal community. Clearly, these shifts should be more pronounced in open canopy gaps created by spring fires, where the soil is more exposed to direct sun radiation. All of the above imply that fire timing can play a major role in shaping the soil fungal community in general and the EMF community in particular, which in turn can determine species-specific plant establishment and growth (Klironomos et al. 2010; Livne-Luzon et al. 2017b), and plant species’ richness (Klironomos 2002). We thus hypothesized that fire season should have a differential effect on the composition of the soil fungal community, shifting the balance between obligate symbiotic EMF, associated with tree roots, and saprotrophic fungi.

Most studies on the post-fire dynamics of the EMF community have been performed in conifer forests (Dove and Hart 2017), located in temperate and boreal areas. In comparison, much less is known about the effect of fire on the EMF community in fire-prone Mediterranean ecosystems. Notably, a few recent studies have brought new attention to fire effects on EMF communities in Mediterranean habitats dominated by *Quercus* or *Cistus* sp. (Buscardo et al. 2015; Buscardo et al. 2010; de Roman and de Miguel 2005; Hernández-Rodríguez et al. 2013), emphasizing the need to explore these more neglected habitats. Our research aimed to study the effects of fire season on the soil fungal community and specifically on the EMF community in a *Cistus* dominated eastern Mediterranean ecosystem. We manipulated fire seasonality using early and late season prescribed burns, and examined the various effects of fire season on the soil- and ectomycorrhizal fungal communities through both sequencing and pine seedling bioassays.

Succession in Mediterranean woodlands often begins with a pioneer stage of *Cistus salviifolius* followed by *Pinus halepensis* colonization (Ne’eman 1997; Sheffer 2012). *Cistus* is considered an early ‘pioneer’ species that increases in density, especially after fire disturbance (Ne’eman and Izhaki 1999). *Pinus halepensis* is a dominant tree species in natural (Liphschitz and Biger 2001) and planted (Osem et al. 2008) forests in Israel, known for its adaptive post-fire regeneration (Ne’eman 1997; Ne’eman et al. 2004). Since at early successional stages *Cistus* shrubs are the main EMF hosts, we hypothesized that *P. halepensis* colonization should be facilitated by the EMF community characterizing *Cistus*. We therefore compared the pine-associated EMF community under *Cistus* shrubs with that of adjacent open canopy gaps using pine bioassays. Doing so allowed us to examine the soil fungal spore bank, essential for the post-fire regeneration of pines in this ecosystem (Glassman et al. 2016b).

## Materials and Methods

### The study area

The study site was located in Har Yaaran in the Judean lowlands of Israel (600 m ASL, Fig. S1). The climate is Mediterranean, with an average annual precipitation of 500-600 mm; between May and October it hardly rains, while solar radiation is very high (Goldreich 2003). The soil is clayish and shallow due to large limestone plates. The vegetation cover is of a Mediterranean shrubland (garrigue), with patches of small multi-stem trees (e.g., *Quercus calliprinos* and *Rhamnus lycioides*), shrubs (1-1.5 m high; e.g., *Pistacia lentiscus*, *Rhamnus lycioide* and *Calicotome villosa*), dwarf-shrubs (≤1 m; e.g., *Cistus salviifolius*, *Cistus creticus* and *Teucrium divaricatum*), and patches of herbaceous vegetation. The main ectomycorrhizal hosts in the study area are (by order of dominance) *C. salviifolius*, *Q. calliprinos* and *C. creticus*. There is an adjacent planted pine (*P. halepensis* and *P. brutia*) forest uphill of the study area, but there were no mature pine trees and only a few pine seedlings were found in the research plots.

### Experimental design

The experimental system consisted of twelve 50×30 m plots, each divided into eight 5×5 m sampling subplots (Fig. S1). Plots were randomly assigned to one of the three following fire treatments (four plots per treatment): 1) spring burns (due to exceptionally late rains, spring burnings were conducted on the 1st of June 2014), 2) autumn burns (11th September 2014), and 3) unburned control plots.

### Soil sampling

Soil samples were collected at four different sampling periods: 1) Pre-fire soil samples (i.e., March 2014), 2) Post-Spring fire (two weeks after the spring burns, i.e., June 2014), 3) Post-Autumn fire (two weeks after the autumn burns, i.e., Oct- 2014), and 4) Post-fires (~1 year after the collection of pre-fire samples, i.e., June 2015). This experimental design and sampling scheme (Fig.1), enabled us to quantify the net effects of spring and autumn burns on the soil fungal community composition, conveying the actual effects of seasonal fires in typical Mediterranean woodlands.

**Figure 1:**
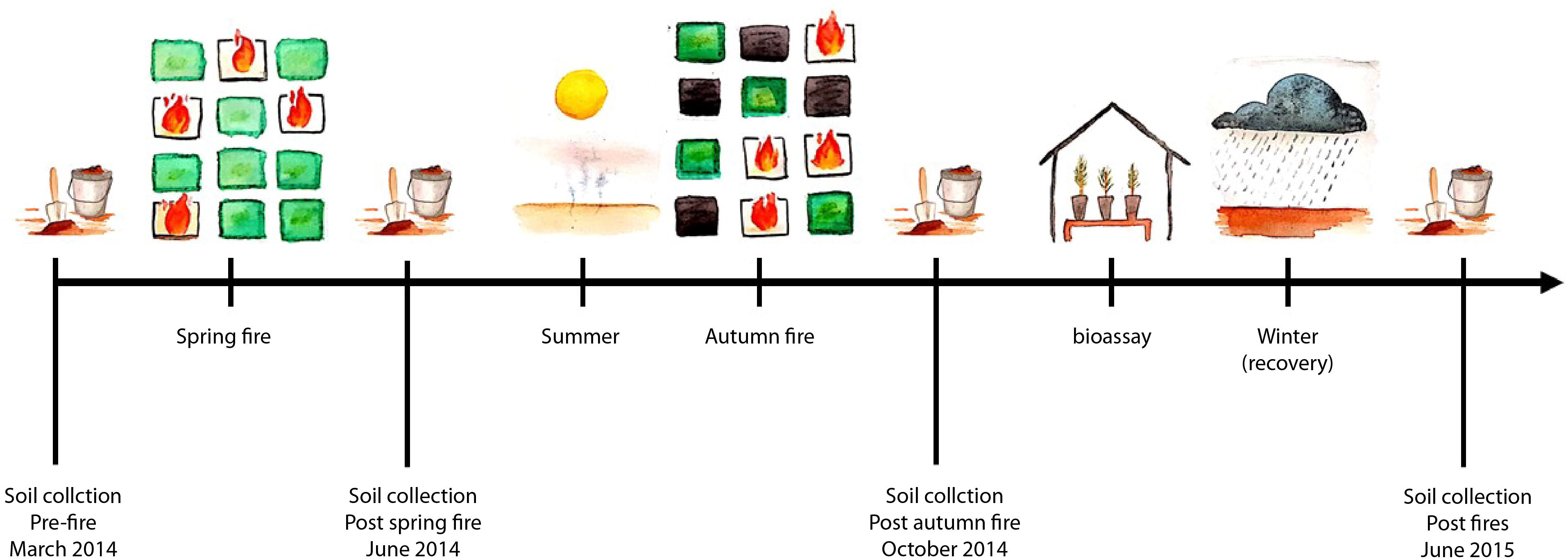
Experimental timescale and sampling scheme. Soil samples were collected from burned and unburned sites before and after spring and autumn burns. Bioassay samples were collected just before the rainy season, reflecting the soil spore bank that germinating plants encounter in the field. This sampling scheme was designed to allow us to quantify the net effects of spring and autumn burns on the soil fungal community composition, conveying the actual effects of seasonal fires in typical Mediterranean woodlands

All samples were collected using the following protocol: three soil cores (10 cm depth, ~ 0.5L) were collected from each of the eight subplots located within each experimental plot. Since the field site is characterized by several rocky patches, each soil core was collected from wherever possible within the 5×5 m subplot, staying within 0.5 m from a *Cistus* shrub (the dominant EMF host in the study area). Each soil core was bagged separately, and all tools were sterilized with 70% ethanol when moving among different subplots to avoid cross-sample contamination. Upon returning to the lab, the three soil cores of each subplot were sieved (2 mm) and homogenized. Then 0.25 g of soil from each sample was directly added to Powersoil DNA tubes (MoBio, Carlsbad, CA USA), and stored in 4°C up to one week before DNA extraction. The remaining soil from the Pre-fire (March 2014) and Post-fires (June 2015) was kept (4°C) in a zip-lock bag for later soil property analysis.

### Greenhouse bioassays

Fungal DNA extracted from the soil may originate from active hyphae or from the soil EMF spore bank (Lindahl et al. 2013; Taylor and Bruns 1999). To assess the inoculation potential of the post-fire EMF community, we bioassayed the soils collected from the study area with *P. halepensis* using standard protocols (Glassman et al. 2016a).

We used the same protocol described above to collect soil samples for the greenhouse bioassay from each of the twelve experimental plots, while distinguishing between two different microhabitats: 1) under a *Cistus* shrub, and 2) an open area without any perennial shrub cover (12 plots × 8 subplots × 2 microhabitats = 192 soil samples). Soil sampling occurred in October 2014, two weeks after the autumn burns and ~4 months after the spring burns, i.e., before the rainy season when most fungi retains activity. Soil samples were air-dried to kill active vegetative fungal hyphae before assaying for resistant propagules (Glassman et al. 2015). *Pinus halepensis* seeds were soaked in water for 48 h, after which they germinated in inert growing media – vermiculite, under controlled conditions in a growth chamber (22 °C, 80% rh, 17 days), and were then planted in the dried soil from each of the 192 bioassay soil samples. Pine seedlings were planted in 200 mL containers using a 1:1 ratio of dried soil and autoclaved sand (121 °C for 20 min ×2), to improve drainage. We controlled for the presence of airborne fungal spores in the greenhouse by adding fifteen replicates of pots containing pine seedlings planted in autoclaved sand. Plants were watered daily and grown in the greenhouse under semi-controlled conditions without fertilizer for approximately six months before harvesting. Treatments were randomized among trays upon initial planting. In total, 207 seedlings were planted (12 plots × 8 subplots × 2 microhabitats = 192 soil samples + 15 controls). After six months, due to harsh summer conditions, only 119 seedlings had survived (8-14 seedlings per plot). Upon harvesting, plants were removed intact from the pots and washed under tap water. Then, roots were thoroughly scanned under a dissecting microscope for colonized root tips. All colonized root tips were removed using sterilized forceps (70% ethanol), inserted into a 1.5 ml Eppendorf tube, and immediately stored in a −20°C freezer. The tubes were immersed in liquid nitrogen at the end of the day, and stored in a −80°C freezer until DNA extraction.

### Molecular identification of species and bioinformatics

Molecular identification of species followed the methods of Glassman et al. (2016b) with minor modifications during the DNA extraction stage. Generally, the ITS1 region was PCR targeted, barcoded and sequenced using Illumina MiSeq technology. For full description of the molecular identification of species and the respective bioinformatic analyses see Supplement S1 and Table S1 in online resource 1. Illumina data were processed using a combination of the UPARSE (Edgar 2013) and QIIME (Caporaso et al. 2010) pipelines following the methods of Smith and Peay (Smith and Peay 2014), and Glassman et al. (Glassman et al. 2016b) with minor modifications related to software updates. Taxonomic assignments were made in QIIME based on the UNITE database (Koljalg et al. 2005). FUNguild was then used to parse OTUs into ecological guilds (Nguyen et al. 2016).

In the greenhouse bioassay, we had ten control pots containing only potting material and plants (no added experimental soil). These root tip samples had low colonization resulting in a total of 48 fungal OTU’s (for all of the controls) with low read abundance (55.08±1.23; mean±1 SE), we thus subtracted these read abundances from the respective data of the bioassay samples. Negative controls from the DNA extraction and PCR stages had all zero reads in them.

### Statistical analyses

Multivariate analyses were performed in PRIMER v.6 of the Plymouth Marine Laboratory (Clarke and Warwick 1994). Relative abundances were fourth-root transformed (Clarke and Warwick 1994; Clarke et al. 2008). A permutational MANOVA (PERMANOVA) based on Bray–Curtis similarity matrix (Anderson et al. 2001) followed by non-metric multi-dimensional scaling (nMDS) ordination was performed to test for the combined effect of fire season (whole plot) and sampling treatments (within plot) on the entire fungal community composition (and on the EMF community) using a split-plot experimental design. A similar analysis was used to examine the combined effects of fire season (whole plot treatment) and microhabitat (*Cistus* vs. Open; within plot treatment) on the fungal community composition associated with pine roots (greenhouse bioassay experiment). We examined the same effects on the relative abundance of each fungal OTU in order to search for specific fungal species that were differentially expressed among these bioassay treatments, the p-values obtained from these tests were than corrected for multiple testing using the false discovery rate correction (Benjamini and Hochberg 1995) implemented in the p.adjust function of the R Stats Package (R Development Core Team 2010). To identify the percentage contribution of different fungal OTU’s to observed differences in community composition, we used a similarity percentages routine (SIMPER) (Anderson et al. 2001). In all cases, qualitative similar results were obtained when a square-root or no transformation were applied, as well as when using a Jaccard similarity matrix (Clarke and Warwick 1994) based on presence/absence data (Clarke and Warwick 1994), so unless otherwise mentioned all results refer to the fourth-root Bray-Curtis similarity matrix. To test for the combined effect of fire season and sampling period on the ratio of saprotrophic to EM fungi ((saprotrophic/EM)/total OUT’s) and on the bioassay OTU richness (and several other diversity indexes), we used split-plot ANOVAs with fire season as the whole-plot factor and sampling period as the within-plot factor. These analyses were performed using STATISTICA v.12 (StatSoft, Inc., Tulsa, OK, USA).

## Results

Fire intensity and severity were largely consistent between spring and autumn burns (Table S2). Notably, the amount of variation in fire intensity and severity among experimental plots was higher during spring than during autumn burns (Fig. S2). For full description of the environmental conditions monitored pre- and during the burns, proxies of fire intensity and severity measured during and post the burns, and analyses of these variables see Supplement S2 in online resource 1. Monthly precipitation, daily precipitation, number of rainy days and ambient temperature during the sampling periods are presented in Fig. S3.

Soil properties including phosphate, nitrite, nitrate, total ammonia-nitrogen, soil organic matter content, and pH were all consistent among fire seasons (Table S3). For full description of these variables and their respective analyses, see Supplement S3 in online resource 1.

### Soil fungal community

#### Time and seasonal fire effects on soil fungal richness and diversity

OTU richness and diversity of soil fungi varied significantly among sampling periods (Tables S4-S6 for the entire soil fungal, EMF and saprotrophic fungal communities, respectively; for the most abundant EMF and saprotrophic fungal taxa see Table S7). The effect of fire season was inconsistent among sampling periods (i.e., Fire season × Sampling period interaction). However, this pattern was significant only when examining the entire fungal community, irrespective of the diversity index used (Tables S4-S6). There were no significant differences in OTU richness and diversity among fire treatments in all sampling periods (March 2014, October 2014 and June 2015), except for the post-spring fire period (June 2014; Fig. S4). Immediately after the spring burns, there was a significant reduction in OTU richness and diversity in burned compared to unburned control plots. Approximately one year after the spring burns these differences diminished (Fig. S4).

#### Time and seasonal fire effects on soil fungal Community composition

Community composition of soil fungi varied significantly by season (PERMANOVA: F_3,25.81_= 12.75, P = 0.0001; Table S8; Fig. S5a), but both fire season (F_2,10.53_ = 0.85, P = 0.621) and the interaction between fire season and sampling period (F_5,23.62_ = 0.67, P = 0.921) were not significant. However, there was higher variation in fungal community composition among plots subjected to spring burns (F_10,249_ = 3.42, P = 0.0001). Similar patterns were observed when examining the EMF and saprotrophic fungal community composition (Tables S9 & S10; Fig. S5).

PERMANOVA pair-wise comparisons (Table 1) indicated that there were no significant differences in community composition of the soil fungi among fire seasons during the pre-fire sampling period (*i.e.,* March 2014), and the same holds true when examining the pre-fire EMF and saprotrophic fungal communities. After spring burns (*i.e.,* June 2014), community composition of the entire and saprotrophic fungal communities varied significantly between the control and spring burned plots (t_11.1_=1.52, p=0.024 and t_11.1_=1.59, p=0.020, for the entire and saprotrophic fungal communities, respectively). However, after autumn burns (*i.e.,* October 2014), these differences disappeared (t_6.14_=1.04, p=0.409), probably due to the large difference in community composition between the control and recently burned autumn plots (t_6.14_=1.67, p=0.006, t_6.14_=1.53, p=0.007 and t_6.14_=1.40, p=0.025, for the entire, EM and saprotrophic fungal communities, respectively). This strong effect of autumn burns on the entire soil fungal community translated into nearly-significant differences between the autumn and spring burned plots (t_6.05_=1.24, p=0.058). In the post-fires sampling period (*i.e.,* June 2015), we observed a significant difference between the control and autumn burned plots (t_6.06_=1.39, p=0.031), but only when examining the entire fungal community. In addition, there was a minor nearly significant difference between the control and spring burned plots (t_6.08_=1.25, p=0.058), but not between autumn and spring burned plots (t_6.1_=1.02, p=0.406).When examining a subset of the data including only the post-fire sampling period (i.e., June 2015), there were significant differences in species composition of soil fungi among the three fire treatments (PERMANOVA: F_2,9.93_= 1.49, P= 0.008; Table S11; Fig. 2a; this effect was weaker when examining subsets of the EM and saprotrophic fungal communities: Tables S12 & S13; Figs. S6). In particular, the soil fungal communities varied between the autumn burned and unburned control plots (PERMANOVA pair-wise contrast:t_6.05_ = 1.38, P= 0.030; Table S14). SIMPER analysis (Fig. 2b) illustrated that numerous OUT’s generated 90% of the dissimilarity among fire treatments and the contribution of each one of them was ~1%. Putative EMF taxa such as *Tuber* and *Inocybe* contributed ~14% to this dissimilarity, while putative saprotrophic fungi generated 22-25% of the dissimilarity among fire treatments. These saprotrophic fungi were attributed to various functional guilds such as wood and dung saprotrophs, but most of these were unidentified to a level allowing for the assessment of their exact function (for the most abundant EMF and SAP taxa in the various soil samples see Table S7). Nevertheless, the ratio of saprotrophic to EM fungi ((saprotrophic/EMF)/total OUT’s) varied among both sampling periods (split plot ANOVA: F_3,22_=41.58 p<0.001; Table S15; Fig.3) and fire season treatments (F_2,10_=8.26 p=0.008). Prior to the burns there were no significant differences between experimental plots (t=0.39, p=0.415). Spring burns led to a reduction in the ratio of saprotrophic fungi, but this pattern was not significant (t=−0.64, p=0.520). Autumn burns led to a significant reduction in the ratio of saprotrophic fungi compared to that of unburned control plots (t=3.12, p=0.002), and this reduction resulted in a significant difference between the autumn and spring burned plots (t= 2.66, p=0.008). Approximately one year after the spring burns (i.e., June 2015), the differences among the control and burned plots disappeared (t=1.12, p=0.259). However, there were still significant differences between the autumn and spring burned plots (t=2.14, p=0.032).

**Figure 2:**
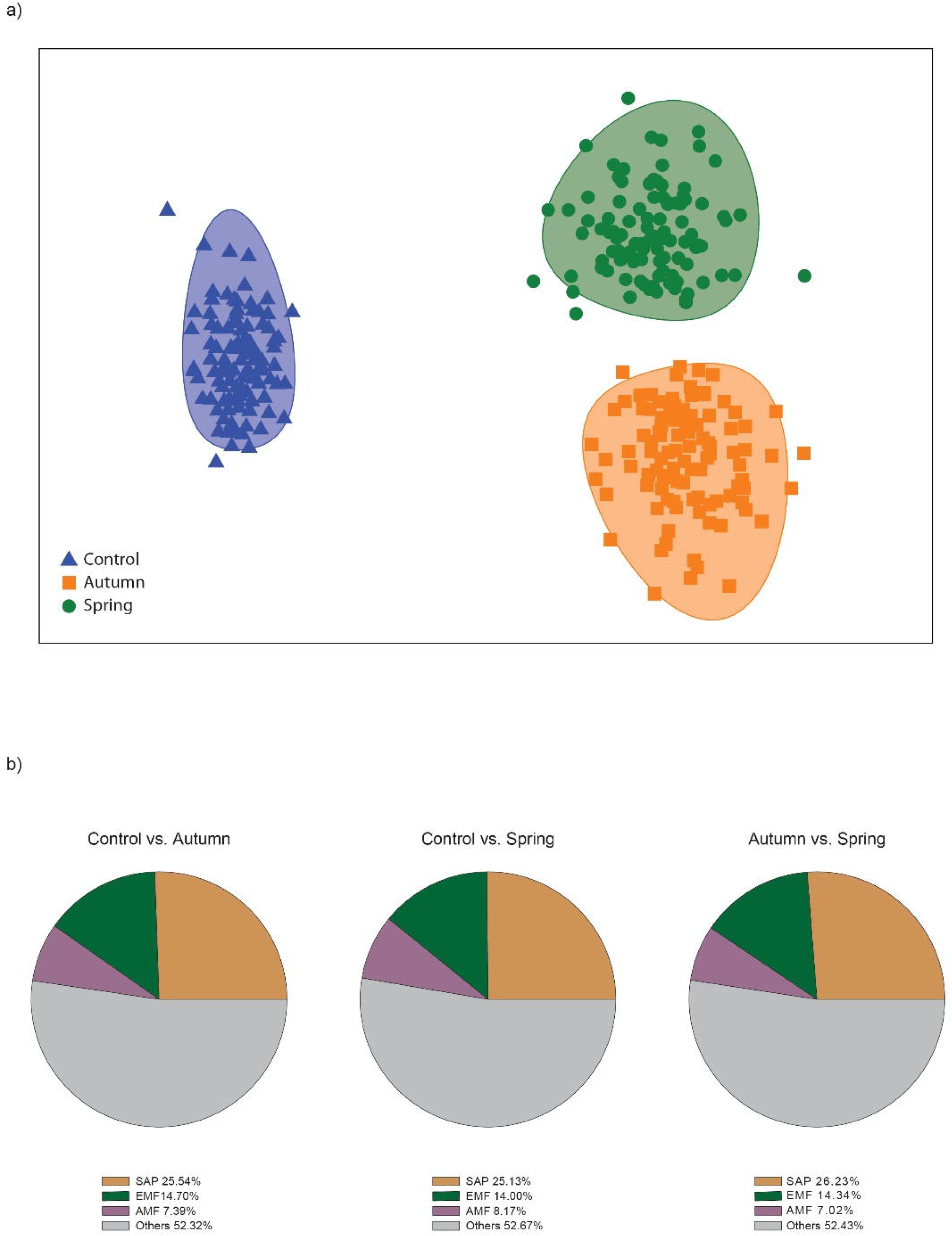
a) Non-metric multi-dimensional scaling (nMDS) ordination with bootstrap of fire season averages, illustrating that soil fungi detected in the post-fire sampling period (i.e., June 2015) vary significantly among fire treatments. Circles represent 95% CI. b) Results of SIMPER analysis illustrating the relative contribution of each of the major fungal functional groups (saprotrophic, EM and AM fungi) to the dissimilarity (Bray-Curtis) among fire treatments. Data includes unassigned fungal sequences and other fungal sequence assignments (e.g., plant pathogen, animal pathogen endophytes etc., each accounted for less than 3% of the dissimilarity).

**Figure 3:**
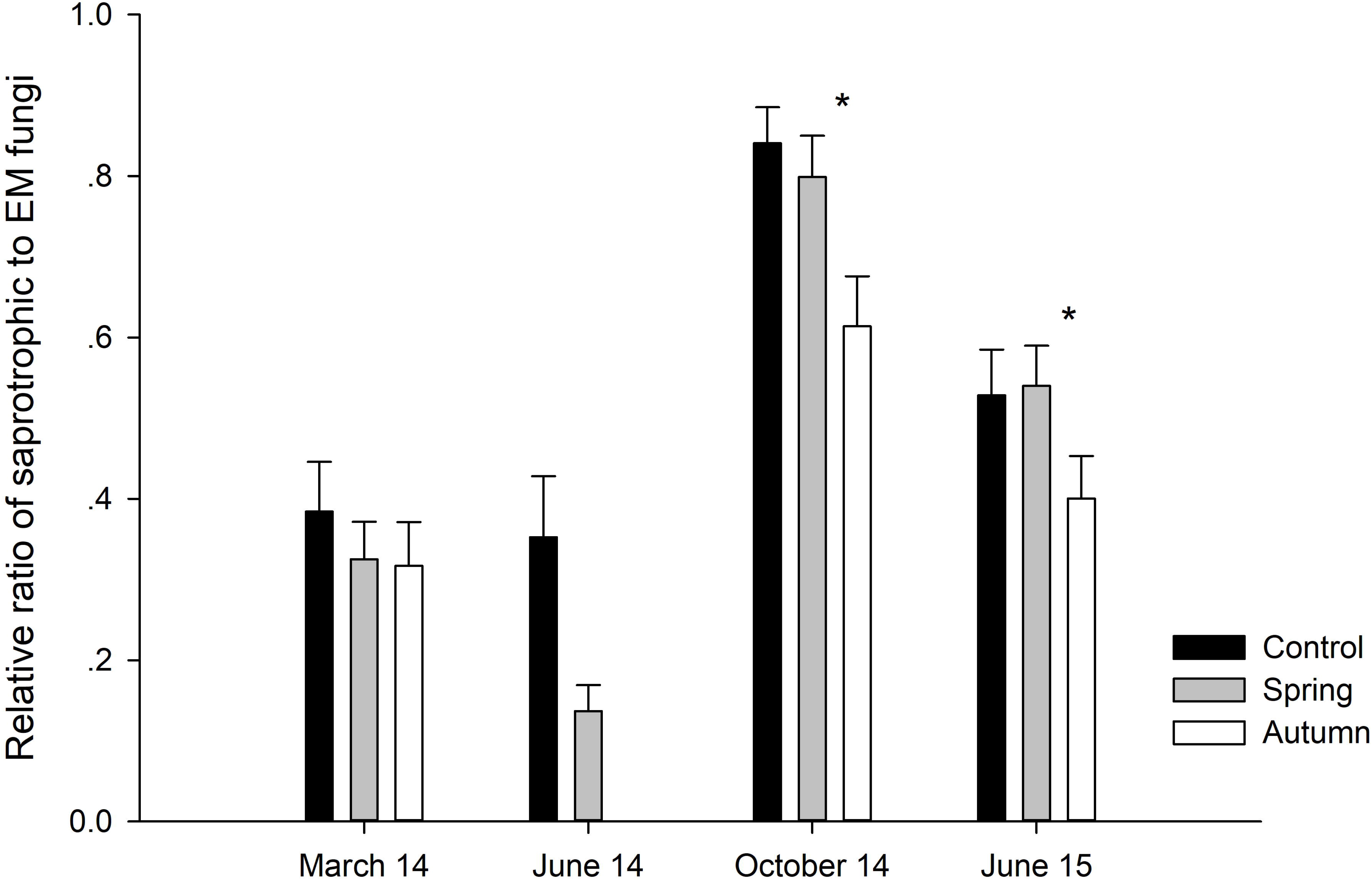
The ratio between OUT’s identified as putative saprotrophic to putative EM fungi out of all OUT’s identified ((saprotrophic/EMF)/total OUT’s) in each sample. * denote significant differences (p<0.05) among the autumn and spring burns within sampling periods.

### Greenhouse bioassay EMF community

Consistent with the results of the soil EMF community, OTU richness and diversity did not vary among fire treatments, nor between microhabitats (Table S16).

Community composition of the bioassay samples did not vary among fire treatments (PERMANOVA: F_2,9_= 0.84, P = 0.605; Table S16; Fig. S6c). Also, there was no significant effect of microhabitat (PERMANOVA: F_1,9_ = 1.76, P = 0.138), nor was there a significant fire treatment by microhabitat interaction (PERMANOVA: F_2,9_ = 0.45, P = 0.916).

### Description of the bioassay EMF community

Regardless of fire season and microhabitat, pine roots in the bioassay were dominated by three major fungal species: *Tuber oligospermum* (21-58%), *Tomentella sp.1* (27-51%) and *Suillus collintus* sp.1 (10-27%). Other fungal species belonging to EMF genera (e.g. other *Tuber spp*., *Inocybe* etc.) accounted for 1-24% of the EMF community (Fig. 4a). These three dominant species were differentially abundant between the *Cistus* and the open microhabitat: *Tuber oligospermum* had higher abundance under *Cistus* shrubs than in the open gaps, while *S. collintus* and *Terfezia* had higher abundance in the open microhabitat (Fig. 4b). Differential expression analysis demonstrated that this pattern was only significant (p<0.01) for *Tuber oligospermum*.

**Figure 4:**
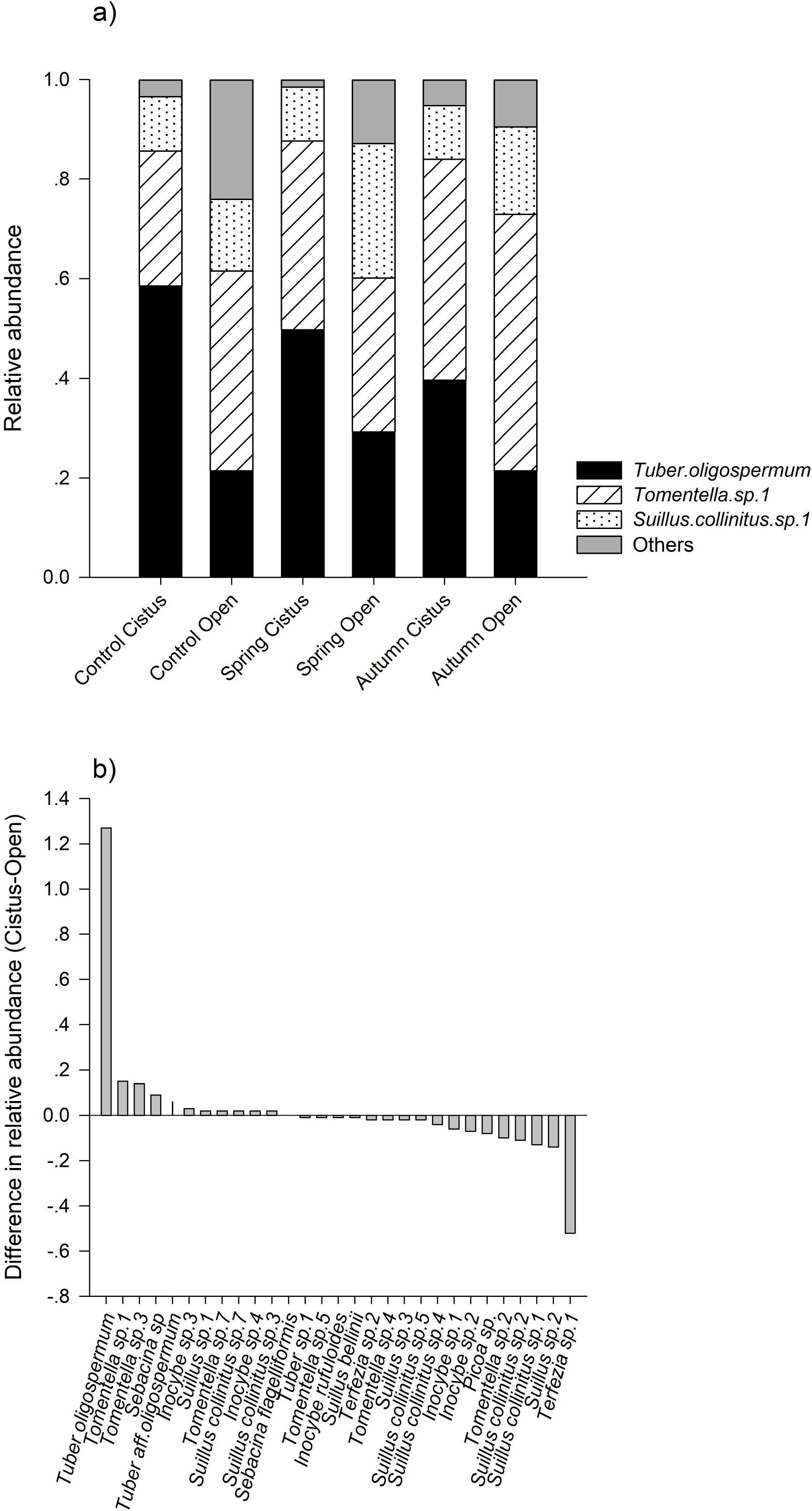
(**a**) Relative sequence abundance of the root-associated EMF species in each of the six treatment combinations of fire treatment and microhabitat. **(b)** Differences in the relative read abundance of the various root-associated EMF species between the two microhabitats (underneath *Cistus* shrubs and open gaps). Positive values are indicative for higher sequence abundance in the *Cistus* microhabitat and negative values indicate higher abundance in open gaps.

## Discussion

We report here the results of the first comprehensive field experiment quantifying the effects of seasonal fires on the soil fungal community in the eastern Mediterranean basin. Fire season caused differential effects on the community composition of soil fungi (Fig.2), driven by alterations within the saprotrophic fungal community (Fig. 3), with the EMF community demonstrating high resilience.

### Differential fire season effects on soil fungi

Our spring and autumn burns did not differ in their intensity and severity, probably due to the specific environmental conditions required for prescribed burns (Allen et al. 1968). Yet, they led to differential modifications in the soil fungal community. Specifically, spring burns caused reductions in soil OTU richness and diversity. But, ~one year after the burns these differences disappeared. Moreover, fire season induced changes in the community composition of soil fungi, which were mostly driven by alterations within the saprotrophic fungal guild (Fig. 2, Fig. 3).

The most parsimonious explanation for the variation in fungal community composition between burning seasons is the time passed since the fires (i.e., areas subjected to spring burns had a longer time to recover before the soil sampling). Another possible explanation is related to fire timing effects. Specifically, seasonal fires occur at different phenological stages of the fungi, potentially resulting in a differential effect on their community composition. For example, fire occurring during the fruiting season might inflict a greater damage than fires occurring during dormancy. Nevertheless, variation in fire intensity and severity among our experimental plots was higher during spring than during autumn burns. Correspondingly, also the variation in community composition of soil fungi was higher during spring, suggesting that fire intensity or severity, regardless of fire season, can lead to changes in the soil fungal community.

Experiencing no significant change due to disturbance (i.e., resistance) and being capable of returning to their pre-disturbance composition (i.e., resilience) are two important features of healthy ecosystems (Shade et al. 2012). Similar to findings from high severity wildfires in CA conifer forests (Glassman et al. 2016b), and in various conifer wildfires in the west-Mediterranean basin (Buscardo et al. 2012; Buscardo et al. 2015; Buscardo et al. 2010), the EMF community represented by both soil samples and pine seedling bioassays appeared to be both resistant and resilient to seasonal fire effects. First, we could not detect a significant effect of fire on EMF richness or diversity. This result is in contrast to most studies of prescribed burns, illustrating a negative effect of fire on EMF richness (Taudière et al. 2017). Second, autumn burns affected EMF community composition, but these differences between the control and autumn burned plots faded quickly and disappeared by the next sampling period (Table 1). Numerous studies have demonstrated that prescribed burns induce changes in the soil fungal community composition (Anderson et al. 2007; Bastias et al. 2006; Hernández-Rodríguez et al. 2015). However, we could not detect a significant effect of fire season on the EMF community composition, neither in soil samples, nor in pine seedling root tips examined in the bioassay experiment (Fig. 2), suggesting high resilience of the EMF spore bank community to fires. This was somewhat surprising, since we are aware of only one other study that did not detect a significant effect of prescribed burns on the soil EMF community composition or richness (Southworth et al. 2011). However, examining the EMF spore bank community after a high-intensity wildfire, Glassman *et al.* (2016b) also demonstrated high resilience of the EMF community in greenhouse bioassays. The ability of the EMF community to survive fire perturbations should contribute to ecosystem stability, since changes in the EMF community can result in structural and functional changes in the respective plant community (Bever et al. 2010).

### Temporal shift in soil fungi

Fungal seasonality has recently been identified as a key feature of natural fungal communities (Averill et al. 2019). Notably, our field experiment lends support to this idea. While examining the soil fungal community, we observed high temporal variation in the soil fungi among sampling periods regardless of fire treatment (i.e., differences appeared also in the unburned control plots). Such variation among sampling periods suggest that fires occurring at different seasons are impacting different pre-fire fungal communities.

Furthermore, EM and saprotrophic fungi compete for limiting resources held within the soil organic matter (Gadgil and Gadgil 1975). These groups have complementary roles in the cycling of nutrients through soil organic matter (Talbot et al. 2014). The observed temporal differences in community composition were mostly related to changes in the relative abundance of saprotrophic fungi among sampling periods (Fig. 2b, Fig.3), which can be related to temporal variation in precipitation attributes. Even though there were no differences in the amount of rain, the number of rainy days marginally varied among sampling periods and was lowest prior to autumn burns (0.8±0.89, mean±1SE; Fig. S3). Conversely, Bell *et al.* (Bell et al. 2009) showed that the saprophytic community of a desert grassland was unaffected by precipitation frequency, however, they suggested that soil temperature, rather than soil moisture strongly influenced fungal carbon use and community structure, and function dynamics. This makes sense, since temperature (i.e., evapotranspiration) affects how much of the soil water will remain available for both plants and fungi. As expected, ambient temperature was higher before autumn burns (25.00±0.91 before autumn vs. 17.35±1.64 before spring burns, mean±1SE). Even though soil moisture was only slightly lower during autumn (3.3±0.22, mean±1SE) than during spring burns (7.69±0.39, mean±1SE), plant water content during autumn burns (0.18±0.01) was half of that measured during spring burns (0.28±0.02, mean±1SE). This suggests that both temperature and precipitation can influence water availability, thus playing an important role in shaping fungal communities in the semi-arid Mediterranean ecosystem.

### The EMF community of Cistus-dominated East-Mediterranean ecosystem

In the eastern Mediterranean ecosystem *Cistus* shrubs are often the main EMF hosts at early successional stages, followed by later successional species such as *Pinus halepensis*. Our data represent the first comprehensive description of the EMF community associated with the understudied *Cistus*-dominated eastern Mediterranean ecosystem. We hypothesized that *P. halepensis* colonization should be facilitated by the EMF community characterizing *Cistus*. We observed high abundance of *Tuber oligospermum* associating with pines grown on soils collected underneath *Cistus* shrubs (Fig. 4), suggesting a newly described link between *T. oligospermum*, *P. halepensis* and the local *Cistus* shrubs (*C. salviifolius*, *C. creticus*). This finding is in concurrence with other studies of *Cistus* dominated ecosystems in the western Mediterranean basin (Comandini et al. 2006), describing the association between *Cistus* and various *Tuber* species. Similarly, but in a conifer-dominated ecosystem, Buscardo *et al.* (2012) and Glassman *et al.* (2016b) found *Rhizopogon* (a Pinaceae specific truffle) increasing in abundance after fire, possibly indicating a pre-adaptation of these hypogeous fungi to fire survival. Since mycorrhizal interactions are often not species-specific (e.g., a mature tree facilitating the establishment of seedlings of a different tree species; Bai et al. 2009; Dickie et al. 2002; Grau et al. 2010; Henry et al. 2015; Kennedy et al. 2003; Kennedy et al. 2012; Richard et al. 2009), these newly described association suggests that interspecific mycorrhizal facilitation is one possible mechanism of facilitation between *Cistus* and *pines*. Such facilitation processes can play an important role in shaping plant community dynamics, vegetation structure and ecosystem functioning (Hayward et al. 2015; Horton et al. 1999). However, the mechanisms governing such facilitation processes are yet to be unravel.

Open canopy gaps were mostly dominated by *Suillus* spores (Fig. 4b). *Suillus* is known for its long-lived (Nguyen et al. 2012), long dispersal distance spores (Glassman et al. 2017; Glassman et al. 2015; Peay et al. 2012) and are known for dominating open microhabitats spore banks. Another dominant fungal genus was *Tomentella*, colonizing pine seedlings grown on soil collected from all fire treatment plots. Similarly, Buscardo *et al*. (Buscardo et al. 2012) reported that *Tomentella ellisii* colonized both pine and oak seedling grown in soil obtained from a short fire return interval site dominated by *Cistus ladanifer*. Interestingly, in a previous study from northern Israel, the genus *Tomentella* also dominated pine seedlings grown in soils collected from a mixed forest site (Livne-Luzon et al. 2017a).

## Conclusions

We observed largest differences due to fire season in the total soil fungal, rather than in the EMF community, and this effect was largely driven by alternations within the saprotrophic fungal guild. Most data on fungal response to fire comes from northwestern USA (Taudière et al. 2017), where fires are typically of higher severity than in the eastern Mediterranean basin. Such fires, as those attained in our study, are less likely to lead to host death, or to inflict direct damage to soil microorganisms. Therefore, Mediterranean fires might induce different selection pressure on the soil biota. Even though the EMF community appeared to be resilient to fire, saprotrophic and EM fungi were documented to compete over similar niche requirements in many ecosystems (Leake et al. 2002). Therefore, these changes in the saprotrophic community composition might have an additional indirect effect on the EMF community composition. Since small-scale changes in carbon inputs can cascade to affect decomposition rates and carbon emissions (Hawlena et al. 2012; Schmitz et al. 2014), these small yet distinct differences in the soil fungal community composition can further affect ecosystem functioning.

## Supporting information

Online resource 1

## Acknowledgements

This research was co-supported by the United States-Israel Binational Science Foundation (BSF Grant 2012081) and Tel-Hai College. We would like to thank the local forester, Mr. Yehezkel Binyamini, and the professional crew of the Jewish National Fund (JNF; Israel’s forest service) who managed the prescribed burnings. We would like to thank Tal Shay and Hadas Ner-Gaon for providing technical support required to install software and packages on the computer cluster and for their help in debugging the pipeline scripts. We also wish to thank Judy Chung and Constantine Klimovitz for their technical help with the plant harvesting and setting-up the sequencing library, and Uri Yogev for his help with the soil properties analyses.

