## Supplementary material for "High resilience of the mycorrhizal community to prescribed seasonal burnings in a Mediterranean woodland": Online resource 1

The following Supporting Information is available for this article:

**Fig. S1** The study site & experimental design.

**Fig. S2** PCA of fire characteristics.

**Fig. S3** Meteorological data: (a) Monthly precipitation, (b) daily precipitation, (c) number of rainy days and (d) ambient temperature, during the sampling periods.

**Fig. S4** (a) OTU richness and (b) Fisher's alpha diversity index of soil fungi.

**Fig. S5** Non-metric multi-dimensional scaling (nMDS) ordinations of: (a) soil fungi, (b) soil saprotrophic fungi and, (c) soil EMF (for all sampling periods)

**Fig. S6** nMDS ordinations of: (a) soil saprotrophic fungi, (b) EMF (only post-fires, June 2015), and (c) root associated EMF in the bioassay experiment.

**Table S1** Proportion of OTU's identified.

**Table S2** Fire characteristics.

**Table S3** Soil chemical properties.

**Table S4** OTU richness and diversity of soil fungi.

**Table S5** Richness and diversity of soil EMF.

**Table S6** Richness and diversity of soil saprotrophic fungi.

**Table S7** The most abundant taxa in soil samples.

**Table S8** PERMANOVA results of soil fungi.

**Table S9** PERMANOVA results of soil EMF.

**Table S10** PERMANOVA results of soil saprotrophic fungi.

**Table S11** PERMANOVA results of soil fungi (post-fires, June 2015).

**Table S12** PERMANOVA results of soil EMF (post-fires, June 2015).

**Table S13** PERMANOVA results of soil saprotrophic fungi (post-fires, June 2015).

**Table S14** PERMANOVA pair-wise comparisons of soil fungi (post-fires, June 2015).

**Table S15** ANOVA results of the ratio of soil saprotrophic fungi.

**Table S16** Richness and diversity of EMF in the bioassay experiment.

**Table S17** PERMANOVA results of EMF in the bioassay experiment.

**Supplement S1** Molecular identification of species and the respective bioinformatics analyses

**Supplement S2** Environmental conditions pre- and during the burns, proxies of fire intensity and severity, and the respective analyses of these variables.

**Supplement S3** Soil chemical properties and their respective analyses.

**Fig. S1** The study site and experimental design.

The study site in Har Yarran (Black Square on the map) located at the Judea Mountains of Israel. Each of the twelve experimental plots was randomly assigned to one of the following three fire treatments: spring burnings, autumn burnings, and unburned control. Each plot included eight 5×5 m subplots.

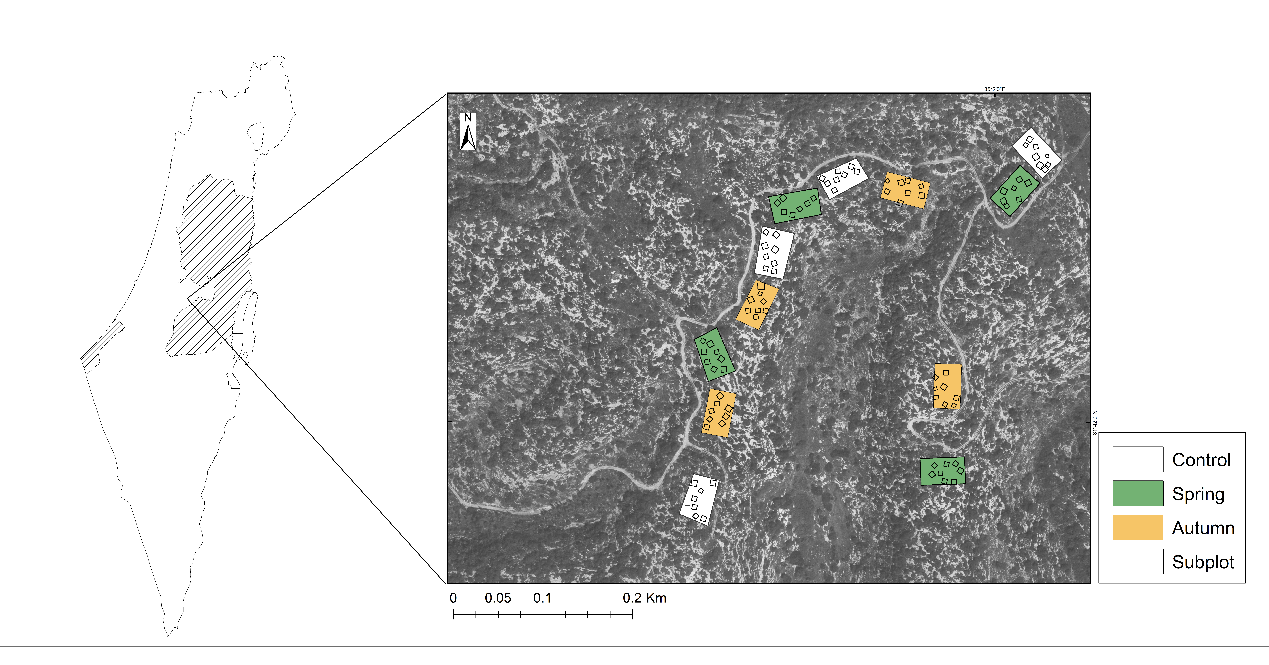

**Fig. S2** Principal Component Analysis (PCA) including measurements taken before, during and after the fires (pre-fire measurements: soil moisture, plant water volume; measurements taken during the fires: wind velocity, relative humidity, flame height; post-fire measurements: percent burned area, burned plant volume).

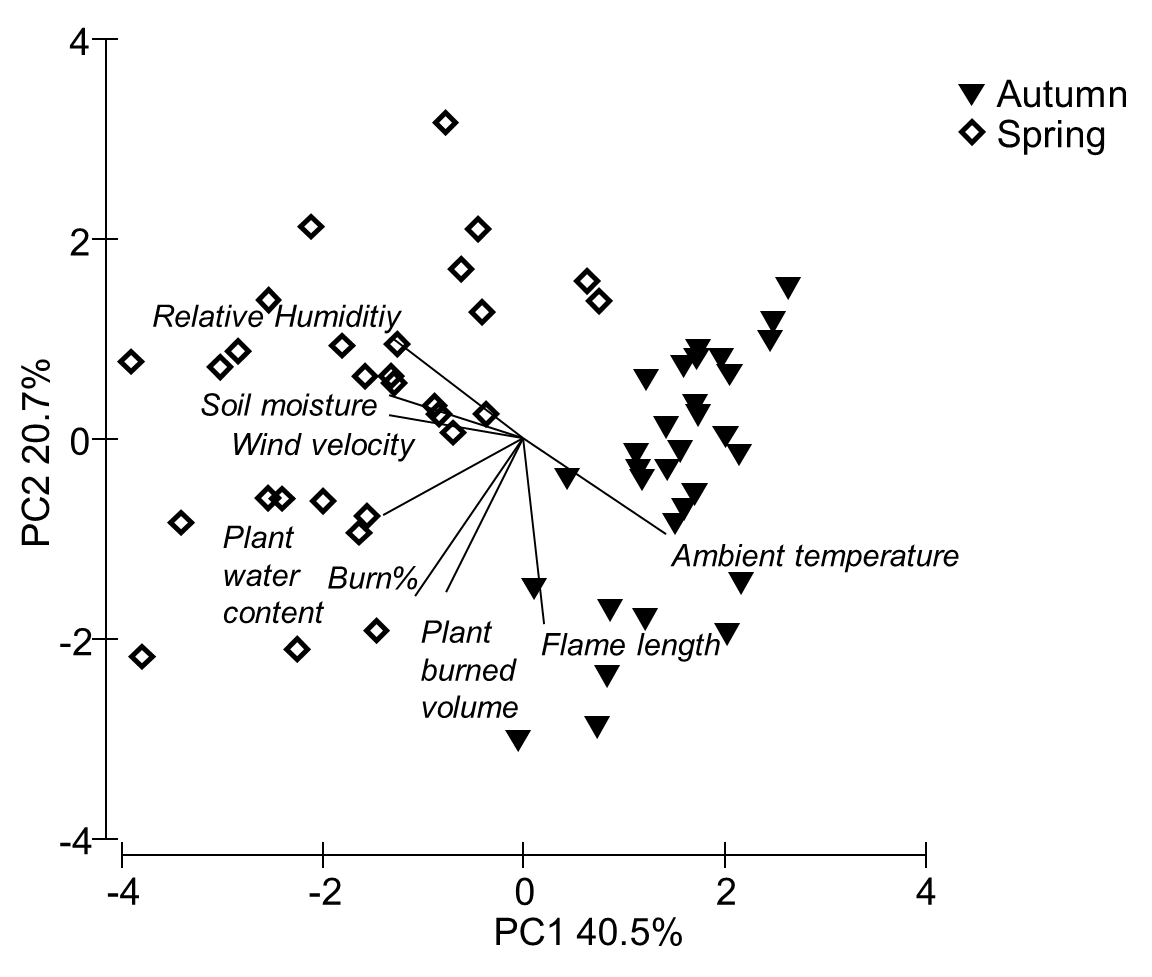

**Fig. S3** Meteorological data: (a) Monthly precipitation, (b) daily precipitation, (c) number of rainy days and (d) ambient temperature, during the sampling periods.

a)

|  | Monthly rain (mm) |
| --- | --- |
| Oct 13- March 14 | 71.6833 |
| June 14- Oct 14 | 5.02 |
| Nov 14- June 15 | 78.7375 |

b)

|  | Maximal daily rain (mm) |
| --- | --- |
| Oct 13- March 14 | 25.2 |
| June 14- Oct 14 | 2.4 |
| Nov 14- June 15 | 29.175 |

c)

|  | Number of rainy days |
| --- | --- |
| Oct 13- March 14 | 4.83333 |
| June 14- Oct 14 | 0.8 |
| Nov 14- June 15 | 6.375 |

d)

|  | Daily temperature (c°) |
| --- | --- |
| Oct 13- March 14 | 17.3545 |
| June 14- Oct 14 | 25.0087 |
| Nov 14- June 15 | 17.3127 |

**Fig S4:** a) OTU richness and b) Fisher's alpha diversity index of soil fungi.

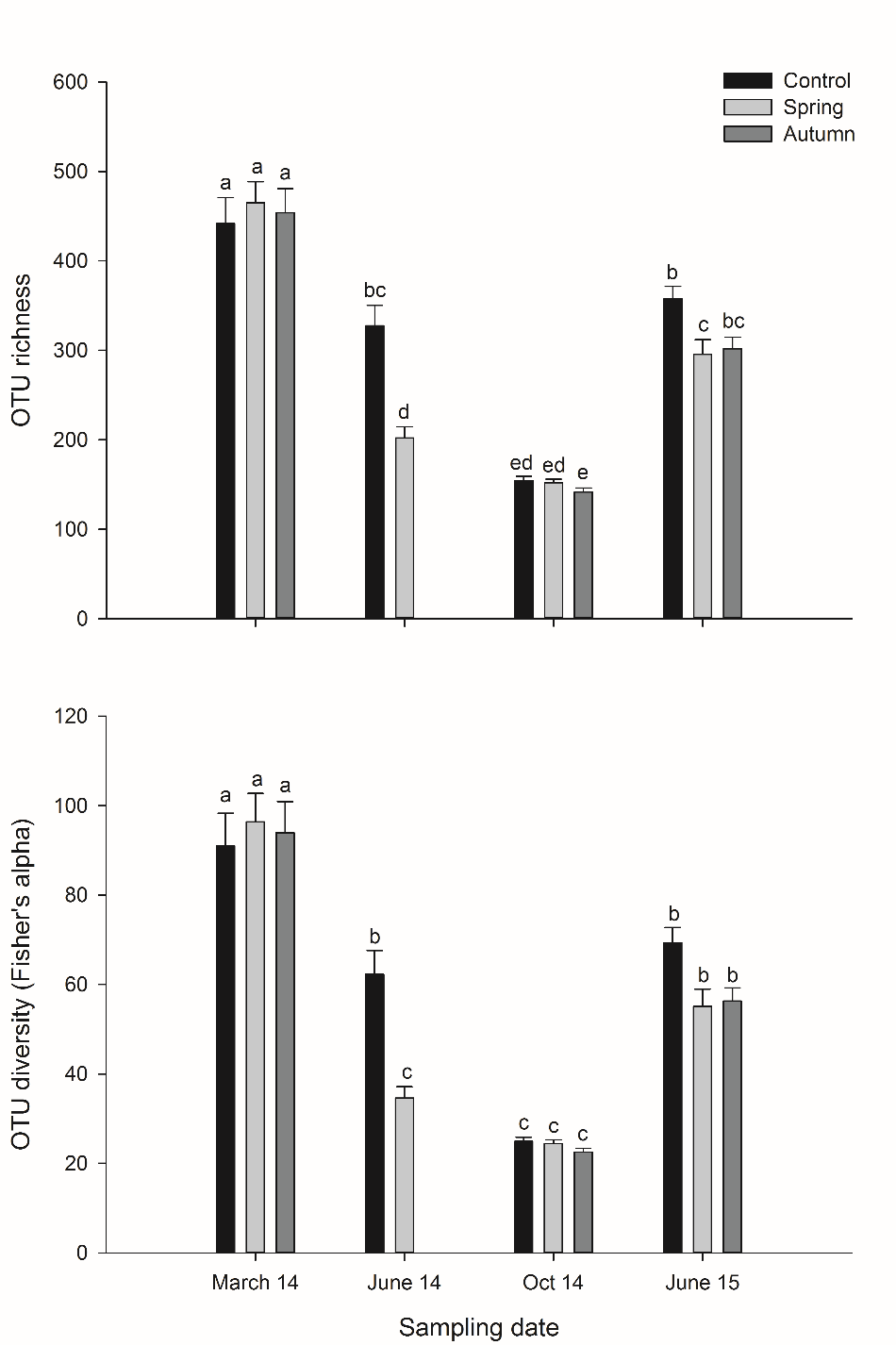

**Fig. S5** Non-metric multi-dimensional scaling (nMDS) ordinations: (a) illustrating that soil fungi community composition vary among sampling periods. While, (b) putative saprotrophic species do not vary among fire treatments nor among sampling periods. However, (c) putative EMF species detected in the soil, vary among sampling periods. Circles represent 95% CI of bootstrap of fire season treatments or the interaction between fire season × sampling period averages.

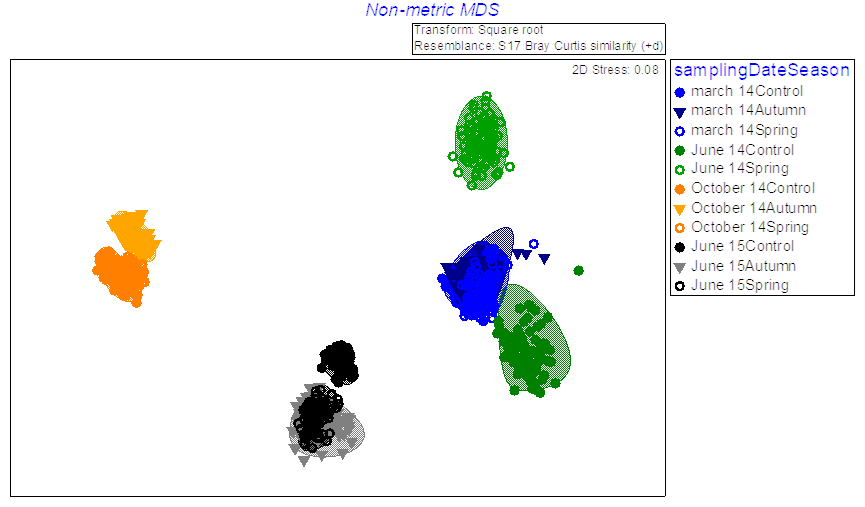

a)

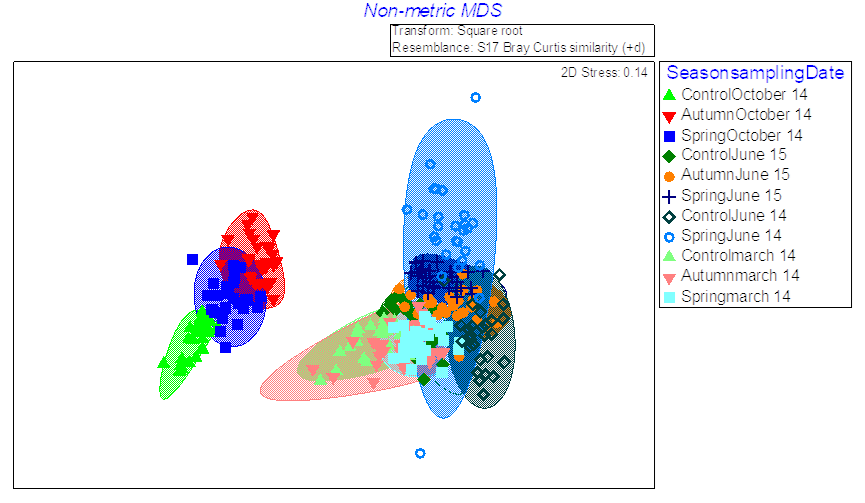

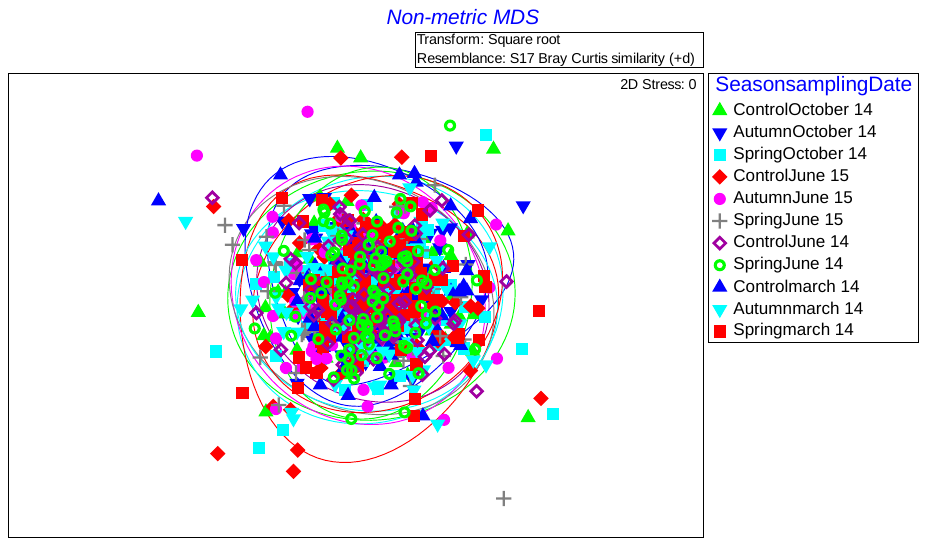

c)

b)

**Fig. S6** Non-metric multi-dimensional scaling (nMDS) ordinations: (a) illustrating that putative saprotrophic species detected in the soil samples during the post-fire sampling period (i.e., June 2015) vary among fire treatments. A similar pattern was obtained when examining the (b) putative EMF species. Nevertheless, the (c) EMF community of the bioassay root-tips do not vary among fire treatments. Circles represent 95% CI of bootstrap of fire season treatments or the interaction between fire season × sampling period averages.

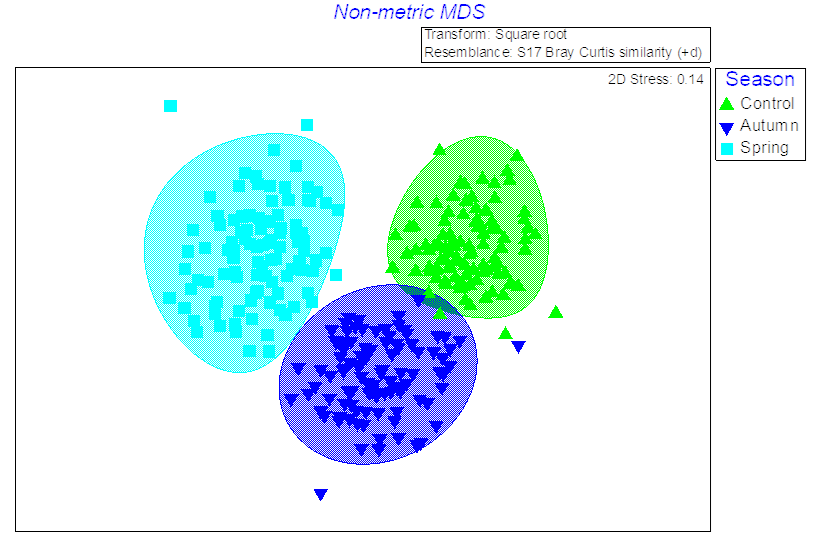

a)

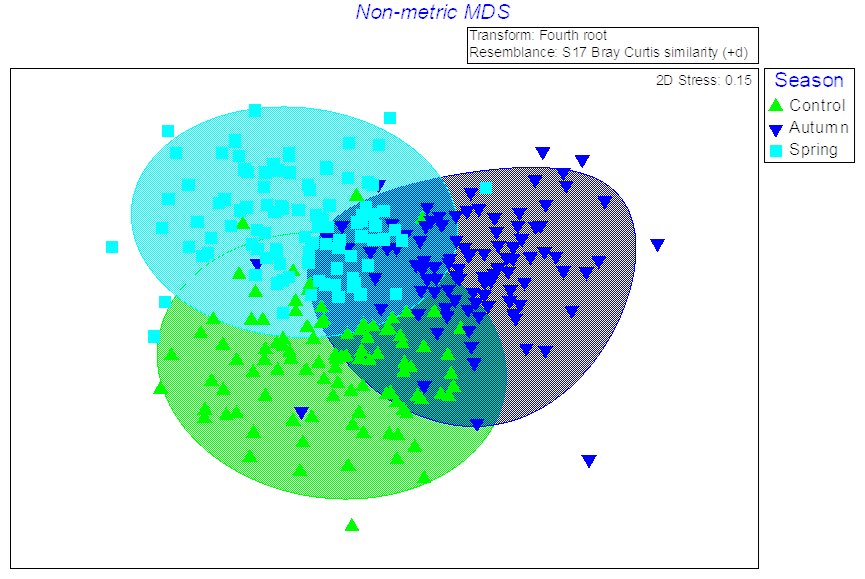

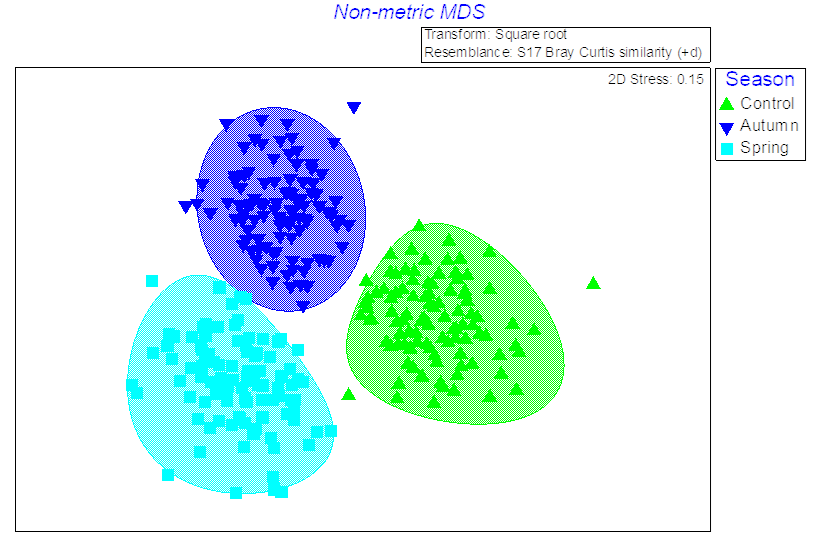
 **Table S1** Proportion of OTU's identified: the number of OTU's that were assigned in BLAST as fungi, and the number of sequences in the total fungal community that were assigned as EMF.

c)

b)

| **Source** | **Total OTU's** | \| **Fungal OTU's** | **Putative EMF** | **FUNguild** |
| --- | --- | --- | --- | --- |
| Soil | 5076 | 4113 | 199 | 838 OTU had a match on FUNguild:  59 highly probable+ probable EMF  130 highly probable+ probable saprotrophic fungi |
| Bioassay | 419 | 298 | 35 | --- |

**Table S2** Fire characteristics: Environmental conditions before (i.e., soil moisture and plant water content) and during the burnings (i.e., ambient temperature, relative humidity and wind velocity). Flame height and direct damage inflicted by the prescribed fires on the vegetation (i.e., proportion of burned area, total burned plant volume). Values represent mean±1SE.

|  | Autumn | Spring | nested ANOVA: fire season |
| --- | --- | --- | --- |
| Soil moisture (%) | 3.3±0.22 | 7.69±0.39 | F_1,52_ = 173.99, *p*<0.001 |
| Plant water content | 0.18±0.01 | 0.28±0.02 | F_1,52_ = 18.62, *p*<0.001 |
| Ambient temperature (˚C) | 29.5±0.19 | 25.68±0.48 | F_1,52_ = 579.3, *p*<0.001 |
| Relative Humidity (%) | 41.05±0.54 | 51.59±1.36 | F_1,52_ = 338.45, *p*<0.001 |
| Wind velocity (mph^-1^) | 3.35±0.24 | 10.48±0.67 | F_1,52_ = 135.06, *p*<0.001 |
| Flame length (cm) | 2.22±0.18 | 1.82±0.16 | F_1,52_=3.30, *p*=0.074 |
| Soil temperature (˚C) | 102±21.03 | 90.94±13.3 | F_1,42_ = 0.298, *p*= 0.587 |
| Burn (%) | 37.11±4.01 | 45.98±4.16 | F_1,52_=2.31,*p*=0.134 |
| Total burned plant volume (m^3^) | 0.32±0.03 | 0.4±0.03 | F_1,52_=3.69,*p*=0.060 |

**Table S3** Results of soil chemical properties (values represent the mean±1SE).

|  | P-PO_4_  (mgN/L)/(g soil) | | N-NO_2_  (mgN/L)/(g soil) | | TAN  (mgN/L)/(g soil) | | N-NO_3_  (mgN/L)/(g soil) | | SOM (g) | | PH | |
| --- | --- | --- | --- | --- | --- | --- | --- | --- | --- | --- | --- | --- |
|  | Pre-fire | Post-fire | Pre-fire | Post-fire | Pre-fire | Post-fire | Pre-fire | Post-fire | Pre-fire | Post-fire | Pre-fire | Post-fire |
| *Control* | 1.56±0.27 | 2.28±0.34 | 2.46±8.42 | 4.16±9.5 | 26.34±2.46 | 52.93±4.16 | 0.27±26.34 | 0.34±52.93 | 0.492±0.054 | 0.498±0.100 | 7.235±0.151 | 7.245±0.149 |
| *Autumn* | 1.96±0.24 | 2.16±0.31 | 3.69±12.11 | 3.3±13.27 | 43.98±3.69 | 61.04±3.3 | 0.24±43.98 | 0.31±61.04 | 0.520±0.019 | 0.671±0.0439 | 7.288±0.130 | 7.252±0.150 |
| *Spring* | 1.87±0.31 | 2.41±0.33 | 3.27±10.84 | 3.51±11.65 | 36.48±3.27 | 52.27±3.51 | 0.31±36.48 | 0.33±52.27 | 0.580±0.062 | 0.620±0.0478 | 7.240±0.114 | 7.383±0.158 |

**Table S4** Results of split-plot analysis of variances (ANOVA), examining the effects of sampling and fire season on the soil fungal OTU richness and several other diversity indexes, with fire season as whole plot factor and sampling as within plot factor. Values represent the mean±1SE. significant differences appear in bold.

|  |  | Richness | 1-Lambda' | Brillouin | d | Fisher | H'(loge) | J' |
| --- | --- | --- | --- | --- | --- | --- | --- | --- |
| Sampling period | Season |  |  |  |  |  |  |  |
| March 14 | Control | 441.73±29.054 | 0.87±0.023 | 3.28±0.159 | 46.76±3.082 | 91.02±7.308 | 3.34±0.162 | 0.55±0.023 |
| March 14 | Spring | 464.87±23.826 | 0.88±0.019 | 3.27±0.159 | 49.21±2.528 | 96.4±6.364 | 3.32±0.161 | 0.54±0.024 |
| March 14 | Autumn | 453.8±26.947 | 0.84±0.027 | 3.08±0.175 | 48.04±2.859 | 93.96±6.971 | 3.14±0.178 | 0.52±0.026 |
| June 14 | Control | 327.67±22.872 | 0.86±0.021 | 2.96±0.162 | 34.66±2.427 | 62.3±5.288 | 3.01±0.165 | 0.52±0.024 |
| June 14 | Spring | 202.17±12.256 | 0.69±0.047 | 2.06±0.159 | 21.34±1.301 | 34.62±2.515 | 2.09±0.16 | 0.4±0.028 |
| October 14 | Control | 154.35±4.913 | 0.75±0.025 | 2.22±0.094 | 16.27±0.522 | 24.93±0.946 | 2.24±0.094 | 0.45±0.017 |
| October 14 | Spring | 151.93±4.206 | 0.75±0.023 | 2.16±0.063 | 16.02±0.447 | 24.44±0.805 | 2.18±0.064 | 0.44±0.012 |
| October 14 | Autumn | 141.6±4.585 | 0.72±0.023 | 2.03±0.08 | 14.92±0.487 | 22.5±0.871 | 2.05±0.08 | 0.42±0.015 |
| June 15 | Control | 357.55±14.119 | 0.91±0.014 | 3.63±0.119 | 37.83±1.498 | 69.3±3.437 | 3.68±0.121 | 0.63±0.018 |
| June 15 | Spring | 295.56±16.531 | 0.94±0.009 | 3.82±0.079 | 31.25±1.754 | 55.1±3.867 | 3.87±0.08 | 0.69±0.013 |
| June 15 | Autumn | 301.8±13.04 | 0.93±0.011 | 3.71±0.1 | 31.91±1.384 | 56.28±2.946 | 3.76±0.102 | 0.66±0.015 |
| Season |  | F_(2,10)_=2.89, p=0.100 | F_(2,10)_=1.82, p=0.210 | F_(2,10)_=1.34, p=0.303 | F_(2,10)_=1.95, p=0.189 | F_(2,10)_=1.50, p=0.266 | F_(2,10)_=1.35, p=0.300 | F_(2,10)_=0.99, p=0.402 |
| Plot(Season) |  | F_(10,22)_=1.13, p=0.380 | F_(10,22)_=1.09, p=0.400 | F_(10,22)_=1.17, p=0.353 | F_(10,22)_=1.14, p=0.370 | F_(10,22)_=1.16, p=0.361 | F_(10,22)_=1.17, p=0.352 | F_(10,22)_=1.14, p=0.374 |
| Sampling period |  | **F_(3,22)_=112.82, p<0.001** | **F_(3,22)_=36.38, p<0.001** | **F_(3,22)_=71.30, p<0.001** | **F_(3,22)_=89.96, p<0.001** | **F_(3,22)_=78.75, p<0.001** | **F_(3,22)_=71.68, p<0.001** | **F_(3,22)_=56.67, p<0.001** |
| Season×Sampling |  | **F_(3,22)_=112.82, p<0.001** | **F_(3,22)_=36.38, p<0.001** | **F_(3,22)_=71.30, p<0.001** | **F_(3,22)_=89.96, p<0.001** | **F_(3,22)_=78.75, p<0.001** | **F_(3,22)_=71.68, p<0.001** | **F_(3,22)_=56.67, p<0.001** |

**Table S5** Results of split-plot analysis of variances (ANOVA), examining the effects of sampling and fire season on the soil EMF richness and several other diversity indexes, with fire season as whole plot factor and sampling as within plot factor. Values represent the mean±1SE. significant differences appear in bold.

|  |  | Richness | 1-Lambda' | Brillouin | d | Fisher | H'(loge) | J' |
| --- | --- | --- | --- | --- | --- | --- | --- | --- |
| Sampling period | Season |  |  |  |  |  |  |  |
| March 14 | Control | 8.04±0.945 | 0.54±0.04 | 1.08±0.083 | 1.35±0.077 | 1.66±0.12 | 0.88±0.109 | 0.49±0.036 |
| March 14 | Spring | 9.17±0.995 | 0.53±0.039 | 1.06±0.077 | 1.51±0.081 | 1.84±0.113 | 0.86±0.104 | 0.45±0.031 |
| March 14 | Autumn | 8.69±1.03 | 0.51±0.043 | 1.03±0.092 | 1.54±0.114 | 1.92±0.168 | 0.79±0.108 | 0.44±0.035 |
| June 14 | Control | 8.5±0.557 | 0.49±0.053 | 0.95±0.097 | 1.15±0.097 | 1.44±0.187 | 0.99±0.109 | 0.48±0.058 |
| June 14 | Spring | 8.17±0.72 | 0.46±0.055 | 0.92±0.11 | 0.98±0.087 | 1.16±0.107 | 0.94±0.111 | 0.47±0.054 |
| October 14 | Control | 2.79±0.395 | 0.48±0.063 | 0.47±0.071 | 0.79±0.105 | 1.82±0.466 | 0.56±0.086 | 0.68±0.064 |
| October 14 | Spring | 3.89±0.464 | 0.46±0.055 | 0.64±0.098 | 0.88±0.098 | 1.39±0.153 | 0.72±0.107 | 0.62±0.052 |
| October 14 | Autumn | 4.63±0.36 | 0.46±0.047 | 0.74±0.073 | 0.94±0.088 | 1.45±0.199 | 0.83±0.088 | 0.58±0.049 |
| June 15 | Control | 6.32±0.472 | 0.46±0.034 | 0.82±0.062 | 0.85±0.064 | 1.06±0.081 | 0.86±0.065 | 0.52±0.04 |
| June 15 | Spring | 6.63±0.481 | 0.48±0.045 | 0.94±0.09 | 0.94±0.074 | 1.16±0.092 | 0.98±0.093 | 0.54±0.04 |
| June 15 | Autumn | 6.9±0.521 | 0.41±0.039 | 0.79±0.076 | 0.93±0.061 | 1.12±0.077 | 0.81±0.078 | 0.44±0.035 |
| Season |  | F_(2,10)_=0.28, p=0.756 | F_(2,10)_=0.85, p=0.455 | F_(2,10)_=0.03, p=0.967 | F_(2,10)_=0.46, p=0.638 | F_(2,10)_=0.08, p=0.917 | F_(2,10)_=0.17, p=0.983 | F_(2,10)_=2.48, p=0.132 |
| Plot(Season) |  | F_(10,22)_=1.12, p=0.309 | F_(10,22)_=1.65, p=0.149 | F_(10,22)_=1.96, p=0.081 | F_(10,22)_=1.19, p=0.341 | F_(10,22)_=1.45, p=0.214 | F_(10,22)_=1.29, p=0.285 | F_(10,22)_=1.45, p=0.213 |
| Sampling period |  | **F_(3,22)_=9.21, p<0.001** | F_(3,22)_=1.398, p=0.262 | **F_(3,22)_=13.43, p<0.001** | **F_(3,22)_=23.03, p<0.001** | **F_(3,22)_=5.51, p=0.003** | F_(3,22)_=1.16, p=0.341 | **F_(3,22)_=8.20, p<0.001** |
| Season×Sampling |  | F_(3,22)_=0.16, p=0.972 | F_(3,22)_=0.27, p=0.922 | F_(3,22)_=1.72, p=0.165 | F_(3,22)_=0.67, p=0.648 | F_(3,22)_=0.41, p=0.831 | F_(3,22)_=0.56, p=0.725 | F_(3,22)_=0.48, p=0.783 |

**Table S6** Results of split-plot analysis of variances (ANOVA), examining the effects of sampling and fire season on the soil saprotrophic fungal richness and several other diversity indexes, with fire season as whole plot factor and sampling as within plot factor. Values represent the mean±1SE. significant differences appear in bold.

|  |  | Richness | 1-Lambda' | Brillouin | d | Fisher | H'(loge) | J' |
| --- | --- | --- | --- | --- | --- | --- | --- | --- |
| Sampling period | Season |  |  |  |  |  |  |  |
| March 14 | Control | 23.39+2.86 | 0.68+0.055 | 1.75+0.158 | 4.56+0.33 | 7.12+0.676 | 1.46+0.199 | 0.56+0.048 |
| March 14 | Spring | 25.31+2.667 | 0.72+0.046 | 1.91+0.131 | 5.13+0.223 | 8.6+0.65 | 1.62+0.194 | 0.6+0.041 |
| March 14 | Autumn | 22.44+2.786 | 0.71+0.049 | 1.8+0.135 | 4.8+0.274 | 7.98+0.648 | 1.46+0.189 | 0.59+0.047 |
| June 14 | Control | 22.75+1.931 | 0.71+0.068 | 1.77+0.177 | 3.66+0.266 | 5.49+0.475 | 1.89+0.19 | 0.62+0.06 |
| June 14 | Spring | 12.33+1.18 | 0.59+0.052 | 1.23+0.128 | 2.33+0.224 | 3.7+0.472 | 1.38+0.145 | 0.56+0.047 |
| October 14 | Control | 9.48+0.691 | 0.48+0.054 | 0.93+0.107 | 1.67+0.127 | 2.44+0.233 | 1.04+0.122 | 0.48+0.054 |
| October 14 | Spring | 8.61+0.762 | 0.39+0.052 | 0.75+0.098 | 1.45+0.158 | 2.23+0.385 | 0.84+0.116 | 0.41+0.052 |
| October 14 | Autumn | 6.37+0.464 | 0.45+0.052 | 0.74+0.081 | 1.24+0.123 | 2.17+0.382 | 0.87+0.1 | 0.49+0.054 |
| June 15 | Control | 23.23+1.036 | 0.7+0.036 | 1.77+0.107 | 3.48+0.172 | 5.01+0.317 | 1.86+0.114 | 0.6+0.035 |
| June 15 | Spring | 17.26+1.342 | 0.71+0.03 | 1.7+0.094 | 2.56+0.216 | 3.5+0.367 | 1.76+0.101 | 0.64+0.032 |
| June 15 | Autumn | 17.69+1.289 | 0.72+0.029 | 1.72+0.089 | 2.73+0.182 | 3.79+0.285 | 1.8+0.093 | 0.65+0.027 |
| Season |  | F_(2,10)_=1.26, p=0.321 | F_(2,10)_=1.14, p=0.357 | F_(2,10)_=2.09, p=0.174 | F_(2,10)_=3.76, p=0.058 | F_(2,10)_=1.24, p=0.329 | F_(2,10)_=0.54, p=0.593 | F_(2,10)_=0.40, p=0.681 |
| Plot(Season) |  | F_(10,23)_=1.23, p=0.321 | F_(10,22)_=0.83, p=0.598 | F_(10,22)_=1.01, p=0.460 | F_(10,22)_=0.90, p=0.546 | F_(10,22)_=0.92, p=0.525 | F_(10,23)_=1.59, p=0.167 | F_(10,22)_=1.12, p=0.378 |
| Sampling period |  | **F_(3,22)_=11.42, p<0.001** | **F_(3,22)_=24.92, p<0.001** | **F_(3,22)_=51.62, p<0.001** | **F_(3,22)_=75.96, p<0.001** | **F_(3,22)_=61.18, p<0.001** | **F_(3,22)_=9.34, p<0.001** | **F_(3,22)_=11.29, p<0.001** |
| Season×Sampling |  | F_(5,22)_=0.45, p=0.805 | F_(5,22)_=1.37, p=0.267 | F_(5,22)_=2.26, p=0.079 | **F_(5,22)_=3.14, p=0.025** | **F_(5,22)_=3.01, p=0.029** | F_(5,22)_=0.28, p=0.914 | F_(5,22)_=1.58, p=0.197 |

**Table S7** The most abundant EMF and saprotrophic fungal taxa (i.e., top 20 of both EMF and Saprotrophic fungi) in soil samples. Data used for these tables includes only highly probable and probable taxonomic assignments by FUNguild.

| Taxon | Total read abundance | Average read abundance | Guild |
| --- | --- | --- | --- |
| Thelephoraceae | 57691 | 5245 | Ectomycorrhizal-Undefined Saprotroph |
| Tuber | 56712 | 5156 | Ectomycorrhizal |
| Tomentella | 53916 | 4901 | Ectomycorrhizal |
| Inocybaceae | 51779 | 4707 | Ectomycorrhizal |
| Sebacinaceae | 44234 | 4021 | Ectomycorrhizal |
| Tricholoma | 37005.5 | 3364 | Ectomycorrhizal-Fungal Parasite |
| Humicola | 30242 | 2749 | Undefined Saprotroph-Wood Saprotroph |
| Thelebolus | 21685 | 1971 | Dung Saprotroph-Endophyte-Undefined Saprotroph |
| Lasiosphaeriaceae | 13289 | 1208 | Undefined Saprotroph |
| Phaeosphaeriaceae | 9715 | 883 | Fungal Parasite-Plant Pathogen-Plant Saprotroph |
| Cenococcum | 8647 | 786 | Ectomycorrhizal |
| Lepiota | 8198.5 | 745 | Soil Saprotroph |
| Pleosporaceae | 7239 | 658 | Endophyte-Lichen Parasite-Plant Pathogen-Undefined Saprotroph |
| Sporormiaceae | 7093 | 645 | Dung Saprotroph-Plant Saprotroph |
| Cladorrhinum | 5884 | 535 | Animal Pathogen-Dung Saprotroph-Endophyte-Plant Saprotroph-Soil Saprotroph-Wood Saprotroph |
| Mycosphaerellaceae | 5106 | 464 | Plant Pathogen-Undefined Saprotroph |
| Hypochnicium | 4344.5 | 395 | Undefined Saprotroph |
| Stemphylium | 4033.5 | 367 | Plant Pathogen-Wood Saprotroph |
| Filobasidium | 3846 | 350 | Undefined Saprotroph |
| Rutstroemiaceae | 2874.5 | 261 | Plant Saprotroph |
| Corynespora | 2680.5 | 244 | Undefined Saprotroph |
| Orbiliaceae | 2552 | 232 | Wood Saprotroph |
| Cortinarius | 2484.5 | 226 | Ectomycorrhizal |
| Crustoderma | 2455 | 223 | Wood Saprotroph |
| Sporormia | 1864.5 | 170 | Dung Saprotroph |
| Vuilleminia | 1849.5 | 168 | Undefined Saprotroph |
| Myrothecium | 1827.5 | 166 | Undefined Saprotroph |
| Cantharellaceae | 1551.5 | 141 | Ectomycorrhizal |
| Thelephora | 1403.5 | 128 | Ectomycorrhizal |
| Geopora | 496.5 | 45 | Ectomycorrhizal |
| Lactarius | 385 | 35 | Ectomycorrhizal |
| Hygrophoraceae | 251.5 | 23 | Ectomycorrhizal-Undefined Saprotroph |
| Melanogaster | 214.5 | 20 | Ectomycorrhizal |
| Boletaceae | 195 | 18 | Ectomycorrhizal-Fungal Parasite-Plant Saprotroph-Wood Saprotroph |
| Xerocomus | 113 | 10 | Ectomycorrhizal |
| Russula | 89.5 | 8 | Ectomycorrhizal |
| Descolea | 68.5 | 6 | Ectomycorrhizal |

**Table S8** Results of split-plot PERMANOVA, examining the effects of sampling and fire season on the soil fungal community composition, with fire season as whole plot factor and sampling as within plot factor.

| Source | df | Den.df | SS | MS | Pseudo-F | P(perm) | Unique perms | Estimates of components of variation | Sq.root | R^2^ |
| --- | --- | --- | --- | --- | --- | --- | --- | --- | --- | --- |
| Season (Se) | 2 | 10.53 | 9527.8 | 4763.9 | 0.85019 | 0.6214 | 9908 | -11.104 | -3.3323 | 0.012 |
| Sampling(Sa) | 3 | 25.81 | 165000 | 54971 | 12.754 | 0.0001 | 9943 | 810.26 | 28.465 | 0.202 |
| pl(Se) | 10 | 249 | 59505 | 5950.5 | 3.4224 | 0.0001 | 9628 | 218.41 | 14.779 | 0.073 |
| Sexsa** | 5 | 23.62 | 15632 | 3126.4 | 0.67656 | 0.9214 | 9896 | -60.933 | -7.806 | 0.019 |
| plot(Se)xsa** | 23 | 249 | 109000 | 4725.4 | 2.7178 | 0.0001 | 9545 | 465.22 | 21.569 | 0.134 |
| Res | 249 |  | 433000 | 1738.7 |  |  |  | 1738.7 | 41.698 | 0.531 |
| Total | 292 |  | 816000 |  |  |  |  |  |  |  |

**Table S9** Results of split-plot PERMANOVA, examining the effects of sampling and fire season on the EMF community composition, with fire season as whole plot factor and sampling as within plot factor.

| Source | df | SS | MS | Pseudo-F | P(perm) | Unique perms | Den.df | Estimates of components of variation | Sq.root | R^2^ |
| --- | --- | --- | --- | --- | --- | --- | --- | --- | --- | --- |
| Season (Se) | 2 | 11284 | 5642.2 | 0.79451 | 0.6599 | 9907 | 10.71 | -19.303 | -4.3935 | 0.011 |
| Sampling(Sa) | 3 | 77156 | 25719 | 5.118 | 0.0001 | 9921 | 27.18 | 330.97 | 18.193 | 0.073 |
| pl(Se) | 10 | 74784 | 7478.4 | 2.5739 | 0.0001 | 9727 | 249 | 237.14 | 15.399 | 0.071 |
| Sexsa** | 5 | 18007 | 3601.5 | 0.68192 | 0.9309 | 9881 | 23.91 | -68.488 | -8.2758 | 0.017 |
| pl(Se)xsa** | 23 | 123000 | 5367.5 | 1.8474 | 0.0001 | 9636 | 249 | 383.5 | 19.583 | 0.117 |
| Res | 249 | 723000 | 2905.4 |  |  |  |  | 2905.4 | 53.902 | 0.689 |
| Total | 292 | 1050000 |  |  |  |  |  |  |  |  |

**Table S10** Results of split-plot PERMANOVA, examining the effects of sampling and fire season on the saprophytic fungal community composition, with fire season as whole plot factor and sampling as within plot factor.

| Source | df | SS | MS | Pseudo-F | P(perm) | Unique perms | Den.df | Estimates of components of variation | Sq.root | R^2^ |
| --- | --- | --- | --- | --- | --- | --- | --- | --- | --- | --- |
| Season (Se) | 2 | 13254 | 6626.8 | 0.99652 | 0.443 | 998 | 10.68 | -0.30615 | -0.55331 | 0.013 |
| Sampling(Sa) | 3 | 102000 | 33971 | 6.8785 | 0.001 | 996 | 26.8 | 464.33 | 21.548 | 0.102 |
| pl(Se) | 10 | 70116 | 7011.6 | 2.6731 | 0.001 | 996 | 249 | 227.58 | 15.086 | 0.070 |
| Sexsa** | 5 | 19130 | 3826 | 0.73315 | 0.88 | 999 | 23.83 | -56.776 | -7.535 | 0.019 |
| pl(Se)xsa** | 23 | 122000 | 5312.7 | 2.0254 | 0.001 | 995 | 249 | 418.96 | 20.469 | 0.122 |
| Res | 249 | 653000 | 2623 |  |  |  |  | 2623 | 51.215 | 0.653 |
| Total | 292 | 1000000 |  |  |  |  |  |  |  |  |

**Table S11** Results of nested PERMANOVA, examining the effects of fire season on the soil fungal community composition post-fires (*i.e*., June 2015), with fire season as whole plot factor.

| Source | df | SS | MS | Pseudo-F | P(perm) | Unique perms | Den.df | Estimates of components of variation | Sq.root | R^2^ |
| --- | --- | --- | --- | --- | --- | --- | --- | --- | --- | --- |
| Season (Se) | 2 | 8543.9 | 4271.9 | 1.4904 | 0.0087 | 9811 | 9.93 | 54.018 | 7.3497 | 0.046 |
| Sub-samples (Su) | 7 | 14857 | 2122.5 | 1.0841 | 0.1171 | 9638 | 54 | 15.829 | 3.9786 | 0.080 |
| pl(Se) | 9 | 26418 | 2935.3 | 1.4993 | 0.0001 | 9554 | 54 | 139.64 | 11.817 | 0.142 |
| Sexsu | 14 | 29120 | 2080 | 1.0624 | 0.0996 | 9461 | 54 | 34.602 | 5.8824 | 0.157 |
| Res | 54 | 106000 | 1957.8 |  |  |  |  | 1957.8 | 44.247 | 0.570 |
| Total | 86 | 186000 |  |  |  |  |  |  |  |  |

**Table S12** Results of nested PERMANOVA, examining the effects of fire season on the EMF community composition post-fires (*i.e*., June 2015), with fire season as whole plot factor.

| Source | df | SS | | MS | | Pseudo-F | | P(perm) | | Unique perms | | Den.df | | Estimates of components of variation | | Sq.root | | R^2^ | |
| --- | --- | --- | --- | --- | --- | --- | --- | --- | --- | --- | --- | --- | --- | --- | --- | --- | --- | --- | --- |
| Season (Se) | | 2 | 6988 | | 3494 | | 0.78873 | | 0.8066 | | 9897 | | 9.96 | | -35.967 | | -5.9972 | | 0.024 |
| Sub-samples (Su) | | 7 | 22919 | | 3274.2 | | 1.0529 | | 0.3293 | | 9765 | | 54 | | 15.815 | | 3.9768 | | 0.079 |
| pl(Se) | | 9 | 40773 | | 4530.3 | | 1.4568 | | 0.0009 | | 9787 | | 54 | | 202.94 | | 14.246 | | 0.140 |
| Sexsu | | 14 | 48524 | | 3466 | | 1.1146 | | 0.1363 | | 9706 | | 54 | | 100.91 | | 10.046 | | 0.167 |
| Res | | 54 | 168000 | | 3109.7 | |  | |  | |  | |  | | 3109.7 | | 55.765 | | 0.577 |
| Total | | 86 | 291000 | |  | |  | |  | |  | |  | |  | |  | |  |

**Table S13** Results of nested PERMANOVA, examining the effects of fire season on the saprophytic fungal community composition post-fires (*i.e*., June 2015), with fire season as whole plot factor.

| Source | | df | SS | MS | Pseudo-F | P(perm) | Unique perms | Den.df | Estimates of components of variation | Sq.root | R^2^ |
| --- | --- | --- | --- | --- | --- | --- | --- | --- | --- | --- | --- |
| Season (Se) | 2 | | 9494.9 | 4747.5 | 1.258 | 0.143 | 997 | 10.05 | 37.422 | 6.1173 | 0.036 |
| Sub-samples (Su) | 7 | | 20351 | 2907.2 | 1.0102 | 0.438 | 995 | 54 | 2.8131 | 1.6772 | 0.077 |
| pl(Se) | | 9 | 34576 | 3841.7 | 1.3349 | 0.001 | 999 | 54 | 137.68 | 11.734 | 0.130 |
| Sexsu | | 14 | 43838 | 3131.3 | 1.088 | 0.112 | 995 | 54 | 71.738 | 8.4699 | 0.165 |
| Res | | 54 | 155000 | 2878 |  |  |  |  | 2878 | 53.647 | 0.585 |
| Total | | 86 | 265000 |  |  |  |  |  |  |  |  |

**Table S14** Results of pair-wise PERMANOVA comparisons, examining the effects of fire season on the soil fungal community composition post-fires (*i.e*., June 2015).

| Groups | t | P(perm) | Unique perms | Den.df |
| --- | --- | --- | --- | --- |
| Control, Autumn | 1.382 | 0.0299 | 3162 | 6.05 |
| Control, Spring | 1.1252 | 0.1066 | 1670 | 7.06 |
| Autumn, Spring | 1.0375 | 0.3373 | 4305 | 7.21 |

**Table S15** Results of split-plot analysis of variances (ANOVA), examining the effects of sampling and fire season on the ratio of saprotrophic fungi, with fire season as whole-plot factor and sampling as within-plot factor.

| Effect | Fixed/random | SS | DF | MS | Error DF | Error MS | F | p |
| --- | --- | --- | --- | --- | --- | --- | --- | --- |
| Season | Fixed | 1.06825 | 2 | 0.53412 | 9.4945 | 0.064592 | 8.2692 | **0.008** |
| plot(Season) | Random | 0.6549 | 10 | 0.06549 | 27.9066 | 0.061261 | 1.069 | 0.416 |
| Sampling date | Fixed | 7.76063 | 3 | 2.58688 | 33.2904 | 0.062211 | 41.582 | **0.000** |
| Sampling date*Season | Fixed | 0.2911 | 5 | 0.05822 | 25.5354 | 0.060767 | 0.9581 | 0.461 |
| plot(Sampling date*Season) | Random | 1.31802 | 22 | 0.05991 | 230 | 0.074472 | 0.8045 | 0.718 |
| Error |  | 17.12853 | 230 | 0.07447 |  |  |  |  |

**Table S16** Results of split-plot analysis of variances (ANOVA), examining the effects of sampling and fire season on the bioassay OTU richness and several other diversity indexes, with fire season as whole-plot factor and sampling as within-plot factor. Values represent the mean±1SD.

|  |  | Richness | 1-Lambda' | Brillouin | d | Fisher | H'(loge) | J' |
| --- | --- | --- | --- | --- | --- | --- | --- | --- |
| Microhabitat | Season |  |  |  |  |  |  |  |
| *Cistus* | Autumn | 59.04±1.07 | 0.32±0.05 | 0.69±0.09 | 6.07±0.11 | 7.88±0.17 | 0.69±0.1 | 0.17±0.02 |
| *Cistus* | Control | 62.67±1.51 | 0.39±0.04 | 0.83±0.07 | 6.45±0.16 | 8.44±0.24 | 0.84±0.07 | 0.2±0.02 |
| *Cistus* | Spring | 57.55±1.16 | 0.26±0.04 | 0.58±0.07 | 5.91±0.12 | 7.65±0.18 | 0.59±0.07 | 0.14±0.02 |
| Open | Autumn | 58.17±1.34 | 0.29±0.05 | 0.62±0.1 | 5.98±0.14 | 7.75±0.21 | 0.63±0.1 | 0.15±0.02 |
| Open | Control | 61.79±1.09 | 0.28±0.05 | 0.62±0.08 | 6.36±0.11 | 8.3±0.17 | 0.63±0.08 | 0.15±0.02 |
| Open | Spring | 61.88±1.91 | 0.37±0.06 | 0.82±0.11 | 6.37±0.2 | 8.32±0.3 | 0.83±0.11 | 0.2±0.03 |
| Microhabitat | | F_(1,9)_=0.50, p=0.492 | F_(1,9)_=0.04, p=0.839 | F_(1,9)_=0.007, p=0.932 | F_(1,9)_=0.50, p=0.492 | F_(1,9)_=0.517, p=0.487 | F_(1,9)_=0.007, p=0.933 | F_(1,9)_=0.01, p=0.916 |
| Season |  | F_(2,9)_=2.92, p=0.10 | F_(2,9)_=0.154, p=0.859 | F_(2,9)_=0.427, p=0.663 | F_(2,9)_=2.92, p=0.10 | F_(2,9)_=2.86, p=0.104 | F_(2,9)_=0.429, p=0.662 | F_(2,9)_=0.40, p=0.680 |

**Table S17** Results of nested PERMANOVA, examining the effects of fire season on the bioassay fungal community composition, with fire season as whole plot factor.

| Source | df | SS | MS | Pseudo-F | P(perm) | Unique perms | Den.df | Estimates of components of variation | Sq.root | R |
| --- | --- | --- | --- | --- | --- | --- | --- | --- | --- | --- |
| Se | 2 | 3623.3 | 1811.7 | 0.84303 | 0.6054 | 9932 | 10.26 | -10.248 | -3.2013 | 0.016 |
| Mi | 1 | 3809.3 | 3809.3 | 1.7594 | 0.1382 | 9945 | 10.47 | 33.717 | 5.8067 | 0.016 |
| Pl(Se) | 9 | 19501.0 | 2166.8 | 1.1307 | 0.2569 | 9846 | 96 | 28.262 | 5.3162 | 0.083 |
| SexMi | 2 | 1964.5 | 982.27 | 0.45305 | 0.916 | 9937 | 10.25 | -72.057 | -8.4886 | 0.008 |
| Pl(Se)xMi | 9 | 19687.0 | 2187.5 | 1.1414 | 0.2437 | 9865 | 96 | 61.186 | 7.8221 | 0.084 |
| Res | 96 | 183970.0 | 1916.4 |  |  |  |  | 1916.4 | 43.777 | 0.788 |
| Total | 119 | 233580.0 |  |  |  |  |  |  |  |  |

**Supplement S1** Molecular identification of species and the respective bioinformatic analyses

Frozen root tips were bead beaten (2 × 30 seconds at 4000 rounds per minute), and DNA was extracted from each root tips sample following a modified version of the QIAGEN (Valencia, CA, USA) DNAeasy Blood and Tissue Kit. The soil samples DNA was extracted following the manufacturer's instructions for Powersoil DNA tubes (MoBio, Carlsbad, CA USA). For both soil and root tips samples, we amplified ITS1, which is a part of the internal transcribed spacer (ITS), the DNA barcode for fungi (47), using custom Illumina sequencing primers (48). The forward primer consisted of the ITS1F primer (49) with the forward Illumina Nextera adapter and a two bp “linker” sequence attached to the 5’ end (48). The individually barcoded reverse primers were composed of the ITS2 primer (50) with the Illumina Nextera adapter, linker sequence, and a 12 bp error-correcting Golay barcode (48). PCRs were cleaned using AMPure magnetic beads (Beckman Coulter Inc., Brea, CA, USA), quantified fluorescently with the Qubit dsDNA HS kit (Life Technologies Inc., Gaithersburg, MD, USA), and pooled at equimolar concentration. Libraries were quality checked for concentration and amplicon size using the Agilent 2100 Bioanalyzer (Agilent Technologies, Santa Clara, CA, USA) at the Functional Genomics Laboratory, University of California, Berkeley, CA, USA. We sequenced three different amplicon pools using Illumina MiSeq technology. The pre-fire (march 2014) and post-spring fire (June 2014) soil samples were sequenced at the DNA Technologies Core at the UC Davis Genome center, while the post-autumn fire (Oct 2014) and post-fires (June 2015) soil samples and the bioassay root tip samples were all sequenced, as two different libraries for soil and root tips data, at the QB3 Vincent J. Coates Genomics Sequencing Laboratory, University of California, Berkeley, CA, USA.

*Bioinformatic analyses*

In order to enable comparisons of the soil samples, sequenced as different sequencing libraries, all bioinformatics steps were performed after pooling together soil data. The root tips data was analyzed separately. Illumina data were processed using a combination of the UPARSE (51) and QIIME (52) pipelines following the methods of Smith and Peay (48), and Glassman *et al.* (19) with minor modifications related to software updates. First, the separate FASTQ files of all samples from each Illumina MiSeq sequencing run were concatenated into two separate files, one for each reading direction. Then, sequences were quality trimmed and reads were paired using USEARCH v. 8.1.1861 i86 Linux 64. Paired reads were discarded if they contained > 0.25 expected errors. Second, high quality sequences were grouped into operational taxonomic units (OTUs) using USEARCH (51) based on 97% similarity (a lower threshold of 95% similarity generated qualitatively similar results). Taxonomic assignments were made in QIIME based on the UNITE database (53) accessed on August 22^nd^, 2016, with the QIIME assign_taxonomy.py script, considering a maximum E-value of 0.01. We then built an OTU table in QIIME; the alignments of all remaining sequences were verified using NCBI BLASTN to make sure the results are valid. (See Table S1 for the proportion of sequences in the total fungal community that were identified as EMF). The data collected was divided into two distinct datasets: (1) the OTU table of the pre- and post-fire soil samples, and (2) the OTU table of the root tips samples. In both cases, we analyzed the data in a hierarchical manner; we first analyzed the OTU table including all fungal OTU's. FUNguild was then used to parse OTUs into ecological guilds (54). The OTU table of the soil samples was rarefied to 12,420 sequences per sample to enable comparisons across samples. The root tips OTU table was rarefied to 14,191 sequences per samples.

In the greenhouse bioassay, we had ten control pots containing only potting material and plants (no added experimental soil). These samples had 48 fungal OTU's with low read abundance (55.08±1.23; mean±1 SE), we thus subtracted these read abundances from the respective data of the bioassay samples. Negative controls from the DNA extraction and PCR stages had all zero reads in them.

**Supplement S2** Environmental conditions pre- and during the burns, proxies of fire intensity and severity, and the respective analyses of these variables

**Methods**

Fire properties were quantified during the prescribed burnings (Tsafrir A. and Ovadia O. unpublished data). Flame length, wind velocity, relative humidity, ambient temperature, and soil temperature were measured during the fires. Flame length was estimated by an observer walking behind the fire line and repeatedly estimating flame length. Wind velocity, ambient temperature and relative humidity were recorded using a mobile weather station every 5 minutes. Soil temperature was measured at 10 cm below the top ground layer using eight thermocouples buried in the soil within two random sampling sub-plots in two of the twelve plots (four thermocouples per plot). Total plant water content was measured by weighing four trimmed twigs of five highly abundant plant species (*Pistacia lentiscus*, *Rhamnus lycioide*, *Cistus salviifolius*, *Cistus creticus*, *Quercus calliprinos* and *Calicotome villosa*) from each plot, before- and after oven drying (60 °C, 48 hours). Plant volume in each subplot was calculated by multiplying the average height of each plant species by its coverage area. The total plant water content in each subplot was calculated as the weighted mean of plant water content of different plant species with their respective volumes as weights.

Soil volumetric moisture content was determined by collecting soil samples (7 cm depth) from each of the 96 subplots during each of the two burning seasons. Soil moisture was calculated by weighing these soil samples before- and after oven drying (105° C, 24 hours). Fire severity (the proportion of burned area) was assessed 10 days after the burnings using the same grid of sample points. At each point, it was determined whether the vegetation was burned by the fire (black), died from the heat (dried out, but not burned - heat shock), or was not damaged by the fire (i.e., remained green).

Environmental data was collected by the Israel meteorological service (from the nearest meteorological station located in Beit Jimal). Data was gathered for three sampling periods: 1) Pre-fires conditions: October 2013 till March 2014, 2) Between-fires conditions: June 2014 till October 2014 and 3) post-fires conditions: November 2014 till June 2015.

Statistical analyses

Conditions before the burnings (i.e., soil moisture and plant water content), and during the burnings (i.e., ambient temperature, wind velocity and relative humidity) and fire intensity and severity (i.e., flame length, soil temperature, and proportion of burned area) were analyzed using one-way nested ANOVAs, with fire season as the explanatory variable. All the above mentioned variables, were used to conduct a Principal Component Analysis (PCA) using PREIMER 6 v.6 (55). To test for variation in fire intensity and severity among the four experimental plots nested within each fire season, we used one-way nested ANOVA on the PC1 scores, with fire season as the explanatory variable.

We examined the effect of sampling period on the monthly and daily precipitation (mm), number of rainy days, and ambient temperature using a set of one-way ANOVAs. Due to violation of the assumptions of normality and homogeneity all count data were square-root transformed ( $y=\sqrt{x+0.5}$ ) and all other continuous data were log transformed ( $y=log(x+1)$ )prior to all analyses.

**Results**

Pre-fire conditions were dryer during autumn burns (Table S2): soil water (F_1,52_ = 173.99, *p*<0.001) and plant water contents (F_1,52_ = 18.62, *p*<0.001) were significantly lower during autumn than during spring burnings. Ambient temperature was lower (F_1,52_ = 579.3, *p*<0.001), relative humidity was higher (F_1,52_ = 338.45, *p*<0.001), and wind velocity was higher (F_1,52_ = 135.06, *p*<0.001) during spring than during autumn burns. There were no significant differences in flame length (i.e., proxy of fire intensity; F_1,52_=3.30, *p*=0.074), nor in soil temperature (i.e., proxy of fire intensity) (F_1,42_ = 0.298, *p*= 0.587) between autumn and spring burns (Supporting Information Table S2). Finally, there were no significant differences in the proportion of burned area (F_1,52_=3.69, *p*=0.060), nor in burned plant volume (i.e., proxies of fire severity, F_1,52_=2.31, *p*=0.134) between the two burning seasons (Table S2). Principal component analysis (PCA), indicated that variables which mostly contributed to the first principal component (PC1) represent the conditions before and during the fire, while variables which mostly contributed to PC2 represent proxies of fire intensity and severity (Fig. S2). While there was a clear discrimination between spring and autumn burnings along PC1 (nested ANOVA on PC1; Season: F_1,52_ = 206.2, *p*<0.001), such discrimination was not evident along PC2. This implies that fire severity did not differ significantly between the two seasons. However, the amount of variation observed in fire intensity and severity among experimental plots was not consistent between fire treatments; the variation among plots subjected to spring burns was higher (nested ANOVA on PC2; Plots (season): F_1,52_ = 3.45, *p*<0.001).

We did not detect a significant difference in monthly precipitation among sampling periods (F_2,16_=2.49, p=0.114, Fig. S3a), nor in maximal daily precipitation (F_2,16_=2.63, p=0.102, Fig. S3b). However, the number of rainy days tended to be lower during the months in between the spring and autumn burns (F_2,16_=3.51, p=0.054, Fig. S3c). Also, Ambient temperature was significantly higher during these between burns period (F_2,16_=7.44, p=0.005, Fig. S3d).

**Supplement S3** Soil chemical properties and their respective analyses

**Methods**

The chemical properties of soil samples collected at each experimental plot (3 fire treatments × 4 experimental plots = 12 samples) and their soil mixtures were analyzed in duplicates, according to standard methods (44) and they included: Phosphate (P-PO_4_), nitrite (NO_2_-N), nitrate (NO_3_- N), total ammonia-nitrogen (TAN), soil organic matter content (SOM) and pH (Table S3).

Statistical analyses

We tested the effects of sampling period and fire season on different soil property measures using a set of two-way split-plot ANOVAs, with fire season as the whole plot factor and sampling period as the within plot factor.

**Results**

P-PO_4_ was consistent among fire seasons (F_2,9_=0.02, p=0.979), and between the pre- and post-fires sampling periods (F_1,9_=2.96, p=0.119, Table S3). Even though NO_2_-N did not vary significantly among fire seasons (F_2,9_=0.07, p=0.925), it was higher in post- than in pre-fire samples (F_1,9_=11.01, p=0.008, Table S3). Similarly, NO_3_-N did not vary significantly among fire seasons (F_2,9_=0.21, p=0.814), but was higher in post-fire samples (F_1,9_=12.04, p=0.006, Table S3). Total ammonia-nitrogen (TAN-N) levels were consistent among fire seasons (F_2,9_=0.42, p=0.664), and between sampling periods (F_1,9_=0.12, p=0.729, Table S3). Soil organic matter (SOM) levels were not affected by fire season (F_2,9_=1.71, p=0.234), nor by sampling period (F_1,9_=1.10, p=0.320, Table S3). pH levels were consistent among fire seasons (F_2,9_=1.3, p=0.309), and between sampling periods (F_1,9_=1.2, p=0.300, Table S3).
